# *INTS6* loss of function disrupts transcriptional regulation in mild intellectual disability

**DOI:** 10.64898/2026.07.17.737701

**Authors:** Nelli Jalkanen, Kalevi Trontti, Antto J. Norppa, Elisa Rahikkala, Panagiotis Lilis, Pau Puigdevall, Leena Ivancic, Nea Niemimaa, Veikko Vuokila, Nour Assaf, Lea Urpa, Mitja Kurki, Eija Hämäläinen, Outi Kuismin, Aarno Palotie, Mikko J. Frilander, Helena Kilpinen, Olli Pietiläinen

## Abstract

Pathogenic variants in genes involved in transcriptional regulation and RNA processing have emerged as points of functional convergence in neurodevelopmental disorders (NDDs), but their specific disease mechanisms remain unknown. By screening 1,562 Finnish extended families from the Northern Finland Intellectual Disability cohort affected by cognitive impairment, we discovered a family with six affected members carrying a heterozygous loss-of-function variant in *INTS6. INTS6 is* a conserved member of the phosphatase module of the Integrator complex, which regulates RNA polymerase II activity, with a reported role in the pathogenesis of NDDs. To determine the variant’s transcriptomic effects, we performed RNA-sequencing of induced pluripotent stem cells (iPSCs) and iPSC-derived neuronal cells from cases and controls, revealing transcriptome-wide splicing defects, with increased intron retention observed in genes involved in translation, cell cycle and RNA processing in variant carriers. CRISPR-Cas9 knock-in iPSCs confirmed that the variant was associated with downregulation of transcription factors and developmental processes in early neuron differentiation. In addition, downregulated genes in variant carrier neurons were enriched for synaptic genes, suggesting effects on neuronal development. These findings highlight the critical role of *INTS6* in transcriptional regulation of human neurodevelopment and reinforce its association with NDDs.

## Introduction

Intellectual disability (ID) is characterised by impairments in cognitive functioning and adaptive behaviours, with diagnosis typically occurring in childhood due to developmental delay^1,2^. ID syndromes are symptomatically heterogenous and can co-occur with symptoms associated with autism spectrum disorders (ASD) and neurological abnormalities^3,4^. The severity of ID varies widely^3^ and individuals with severe or profound forms often need constant support for daily living, while those mildly affected can live independently in society^5^. The prevalence of ID is around 1% in the general population globally ^1,6–8^, with mild ID being more common than severe ID^7,9^.

Genetic factors play a major role in the aetiology of ID and offer a starting point for biological modelling. Past genetic studies in ID have highlighted a high burden of dominant *de novo* variants in genes with roles in transcriptional regulation and RNA processing, including chromatin modifiers and transcription factors^10–14^. Consistent with the clinical heterogeneity of ID, many of the involved genes are shared across neurodevelopmental disorders (NDDs) and psychiatric disorders, ASD and schizophrenia^15^. This suggests that disruption of a single gene regulatory process can lead to diverse clinical presentations. Despite the growing knowledge of genes affected in ID, the biological mechanisms that mediate the pathogenic effects remain largely unknown, especially in mild ID.

Recently, pathogenic variants in several genes encoding for subunits of the Integrator complex have been linked with NDDs ^16–21^, including ID^22^. The Integrator is a multiprotein complex that associates with the C-terminal domain (CTD) of RNA polymerase II (RNAPII)^23,24^ to regulate the transcription of coding^25–32^ and non-coding RNAs^23,24,33–36^. The Integrator complex has two catalytic domains: an endonuclease domain^37^, essential for the 3’-processing of small nuclear RNAs (snRNAs)^23,38^, and a phosphatase domain that regulates RNAPII activity^39,40^. Predicted dominant loss-of-function variants (pLoFs) in *INTS6* (*Integrator Complex Subunit 6*) of the Integrator phosphatase domain were recently found to be associated with NDDs, including ID and ASD^22^. Loss of *INTS6* alters RNAPII activity by interfering with RNAPII-CTD phosphorylation and by impairing synapse development and structural organization of the cerebral cortex in mouse^22^. However, the role of pathogenic variants in *INTS6* in human neuronal function has not yet been systematically studied.

Here we report a novel heterozygous pLoF *INTS6* variant (GRCh38 chr13:51378376:G:A; NM_012141.3: c.1465C>T, p.Arg489*), which is maternally inherited and carried by six siblings diagnosed with predominantly mild ID in a large family from the Northern Finland Intellectual Disability (NFID) cohort. To understand how partial loss of *INTS6* affects human neurons, we established induced pluripotent stem cell (iPSC) lines from variant carriers (N=5), from unaffected family members (N=3) and from external controls (N=5). In addition, we used CRISPR-Cas9 to correct the variant in one carrier line and to introduce the variant (knock-in) into one control line. We performed RNA-sequencing for cases and controls at different stages of neuronal differentiation as iPSCs, neural progenitor cells (NPCs) and mature neurons, and for the CRISPR-edited lines as iPSCs. This identified a transcriptomic signature of RNA processing dysregulation associated with the heterozygous *INTS6* variant.

## Results

### Loss-of-function variant in INTS6 associates with mild intellectual disability

Deleterious variants in the *INTS6* subunit of the Integrator complex (**Fig S1A**) have recently been associated with NDDs including ID^22^. To assess the role of *INTS6* as a cause for mild ID, we leveraged WES data^41^ from the NFID cohort (N=3,974, N=1,772 with ID) to search for pLoF variants in the *INTS6* gene. We identified seven individuals in one large family carrying a protein-truncating variant (PTV) (c.1465C>T, p.Arg489*) in *INTS6* **(Fig 1A),** located in exon 12 (ENSE00003543787) of transcript ENST00000311234.9 (**Fig 1B**). Using Sanger sequencing, we confirmed the presence of the variant in all carriers and its absence in non-carriers **(Fig. 1A).** Six of the carriers, aged 13–28 years, were diagnosed with non-syndromic ID. The carriers presented mild (N=4) to moderate (N=1) ID (ICD-10 F70 and F71, respectively) or a mixed disorder of scholastic skills (ICD-10 F81.3, N=1). All cases had delayed language development and received speech and/or occupational therapy. Their growth was normal and they did not have visual or hearing deficits. Brain magnetic resonance imaging (MRI) and electroencephalography recordings were normal **(Table S1).** The variant was transmitted from the mother to the six affected offspring but was not transmitted to the unaffected siblings. The mother has dyslexia, completed comprehensive school and vocational school and has not been diagnosed with ID. Overall, this supports co-segregation of the variant with the ID phenotype in the family (segregation p = 0.00195), although the undiagnosed carrier mother may indicate reduced penetrance.

**Figure 1:**
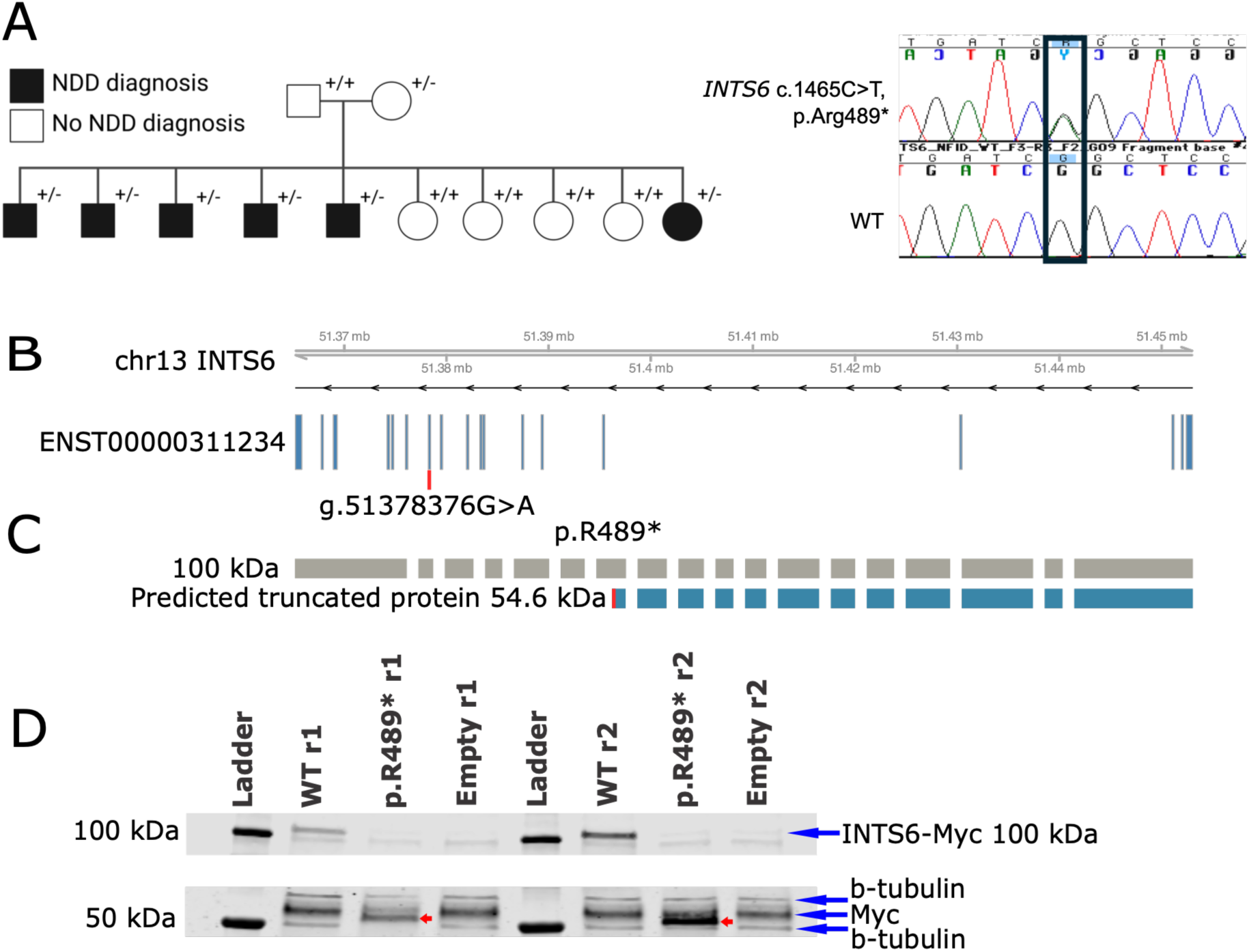
A stop-gain variant in *INTS6* identified in a Finnish extended family with predominantly mild intellectual disability. **(A)** An *INTS6* variant was identified in a family with six non-syndromic ID cases from the NFID study (pedigree, left). The variant was confirmed by Sanger sequencing, demonstrated by a representative Sanger trace (right). **(B)** *INTS6* c.1465C>T, p.Arg489* variant is located in exon 12 of 18 of the canonical MANE select transcript NM_012141.3. **(C)** The chr13:51378376:G:A variant causes a premature stop-codon in *INTS6* (p.R489*) based on UniProt. **(D)** Western blot showing overexpression of the INTS6-Myc fusion protein containing either the control (WT) or mutant (pR489*) allele in HEK293T cells. The overexpression assay was performed in two technical replicates (r1/r2). An empty plasmid was used as the negative control. pR489* samples show a lack of INTS6 protein expression, and the red arrow indicates the 54.6 kDa predicted truncated protein that is expressed in the pR489* samples. The additional band below INTS6-Myc 100 kDa and 54.6 kDa corresponds to endogenous Myc in the cells.

### The INTS6 variant produces a truncated protein consistent with a stop-gain effect

To validate the functional consequences of the pLoF we analysed the protein-level effect of the predicted premature stop codon of the *INTS6* variant. Overexpression of an INTS6-Myc fusion protein containing either the mutant or the reference allele in HEK293T cells confirmed that the mutant allele did not produce the full-length 100 kDa INTS6 protein, consistent with a stop-gain effect **(Fig. 1D).** The pathogenic variant resulted in a truncated, 54.6 kDa INTS6 protein, confirming the predicted functional impact of the variant **(Fig. 1D, Fig. S1B).** To study the biological impact of the partial loss of *INTS6*, we generated iPSCs from 5 cases and 8 controls (**Fig. 2A; iPSC lines described in Table S2**). We first set out to confirm the variant effect in the iPSCs. Consistent with a stop-gain effect, we observed slightly reduced INTS6 protein levels in the case iPSC lines, although this was not statistically significant (N_cases_ = 3, N_controls_ = 3, estimated marginal mean difference = −0.0178, F(1,9) = 4.11, p = 0.0733, nested ANOVA) **(Fig. S1C).**

**Figure 2:**
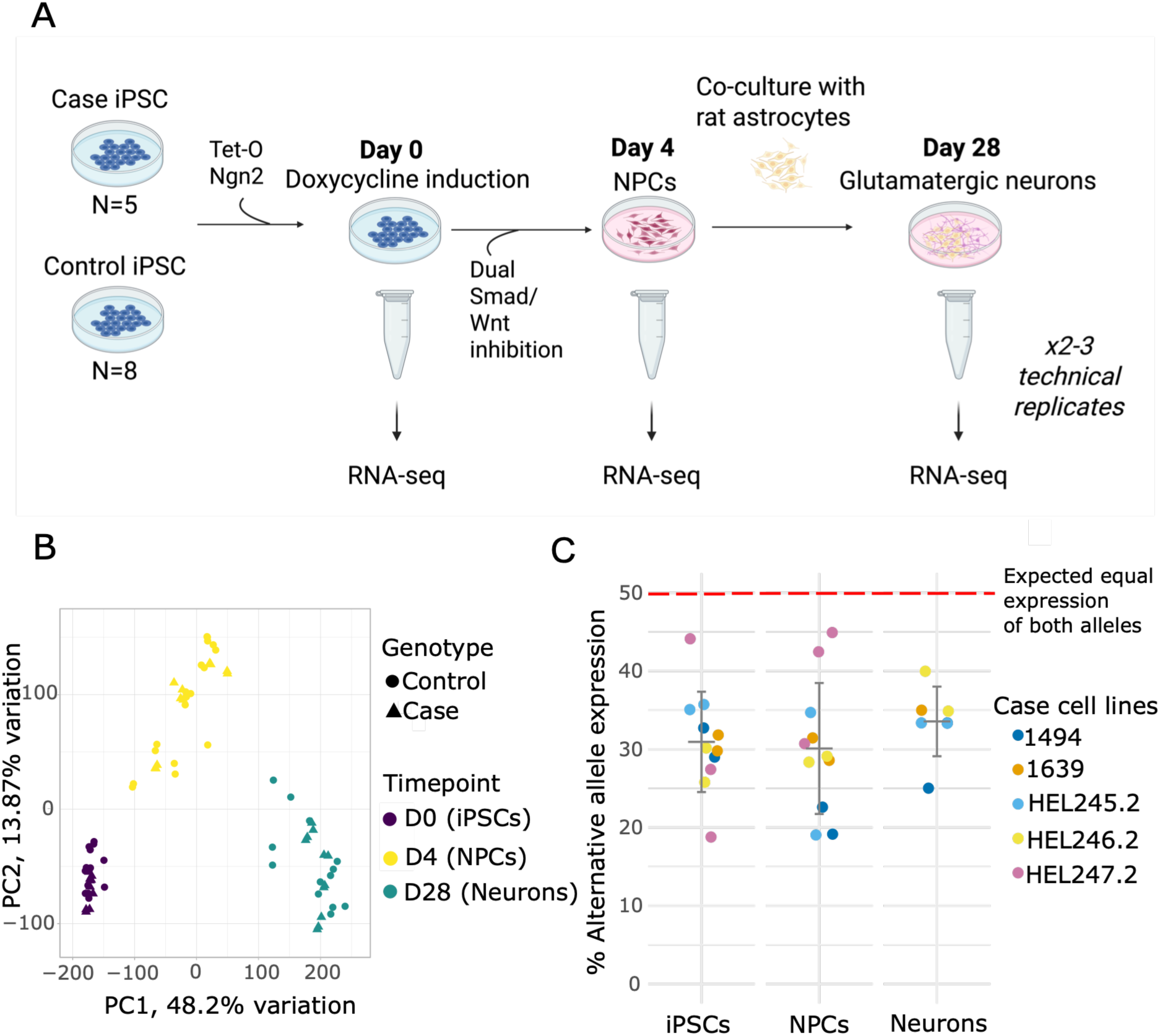
Experimental design and transcriptomic characterisation of neuronal differentiation. **(A)** Schematic of the iPSC-based experimental design to study the transcriptomic consequences of the *INTS6* variant across neuronal differentiation. **(B)** PCA plot of samples that were RNA-sequenced. PC1 loadings correlate with differentiation timepoint (i.e. cell type), while cases and controls cluster together. **(C)** The average allelic expression of the alternative *INTS6* allele (A) in cases is 32.9%, representing a 0.658-fold difference relative to the wild type allele G (p = 7.916 × 10⁻⁹, binomial test).

### iPSC-neuron differentiation is reproducible and comparable between *INTS6* cases and controls

To further study the biological consequences of the *INTS6* variant in neuronal functions, we differentiated the case and control-derived iPSCs towards a glutamatergic neuron fate using *Ngn2* overexpression and dual Smad and Wnt inhibition^42^. From day 4 onwards, neurons were co-cultured with rat astrocytes to promote synapse formation and survival. Prior to differentiation, all iPSC lines were confirmed to have a normal karyotype and to express markers of the undifferentiated state (OCT4, Nanog, and Tra-1-81) using immunocytochemistry **(Fig. S2A)**. We collected samples at three timepoints: D0 (iPSCs), D4 (neuronal progenitor cells; NPCs) and D28 (neurons). NPC and neuron cultures were also characterised with immunocytochemistry, with NPCs expressing PAX6, Nestin and SOX1, but not OCT4, and neurons expressing MAP2 and VGLUT1 **(Fig. S2B-C)**.

To study the transcriptomic consequences of the pathogenic *INTS6* variant across development, we performed RNA-sequencing (RNA-seq) at different timepoints of the differentiation: the baseline (D0, iPSCs), neuronal progenitor timepoint (D4, NPCs), and mature neurons (D28) **(Fig. 2A).** Principal component analysis (PCA) of global gene expression profiles showed separation of samples from different timepoints along the first two principal components (PC1 and PC2), with cases and controls clustering together **(Fig. 2B).** This implicated that differentiations from multiple cell lines were reproducible across the experiment, and that the differentiation capacity of individual samples was not strongly affected by the variant or the differentiation batch **(Fig. S2D).**

We carried out deconvolution of bulk RNA-seq profiles using *Ngn2* D28 neuron single-cell RNA-seq data as a reference^43^ to further assure similarity of cell type composition between cases and controls. The percentage of excitatory neurons among all samples at the D28 timepoint varied between 62.2 - 87.4% (average = 74.3%) **(Fig. S3A)**. We observed no significant differences in the abundance of excitatory neurons between cases and controls (72.7% vs 75.4%, p = 0.27, Wilcoxon test) **(Fig. S3B)**. Similarly, the proportion of NPCs at the D28 timepoint was not significantly different between cases and controls (p = 0.29, Wilcoxon test) and neither was the proportion of peripheral nervous system (PNS) neurons (p = 0.61, Wilcoxon test), suggesting that the differentiation outcome was not influenced by the variant.

### Developmental and synaptic gene programs are downregulated in *INTS6* cases

We next analysed how the *INTS6* variant influenced gene expression during neuronal differentiation. While total mRNA levels of *INTS6* did not show significant differences between cases and controls **(Fig. S2E)**, the mutant allele showed reduced relative expression, with the A allele expressed at 33% versus the expected 50% (0.658-fold relative to the wild type G allele, p = 7.9 × 10⁻⁹, binomial test) **(Fig. 2C)**. We next analysed differential gene expression between cases and controls across the differentiation timepoints. The analysis revealed 687 significantly upregulated genes and 521 significantly downregulated genes in case iPSCs, 109 significantly upregulated and 41 significantly downregulated genes in case NPCs, and 223 significantly upregulated and 111 significantly downregulated genes in case neurons **(Tables S4-S6)**. In all differentiation timepoints, there were more upregulated than downregulated differentially expressed genes (DEGs) in cases: D0 (56.9% upregulated, binomial test p=9.95×10^−7^), D4 (72.7% upregulated, p = 1.31×10^−8^), and D28 (66.8% upregulated, p=4.38×10^−10^), suggesting a consistent effect toward upregulation in case cells.

The DEGs were predominantly timepoint-specific with little overlap of significant genes. This was demonstrated by the 630 upregulated (91.7% of total D0 upregulated DEGs) and 495 downregulated (95.0%) genes specific to D0, 60 upregulated (55.0%) and 23 downregulated (56.1%) specific to D4 and 204 upregulated (91.5%) and 96 downregulated (86.5%) genes specific to D28, with only 42 upregulated and 14 downregulated genes shared between D0 and D4, and 4 upregulated and 3 downregulated genes between D4 and D28 **(Fig. 3A)**. This may suggest that the variant could influence different biological functions at different developmental stages. The overall lower number of DEGs in D4 and D28 samples could be partly explained by the average library size being nearly halved compared to D0 due to the co-culture conditions (72.6 million and 73.3 million reads vs. 132.5 million reads, respectively), which inherently reduces the statistical power to detect low-abundance transcripts **(Table S3).**

**Figure 3:**
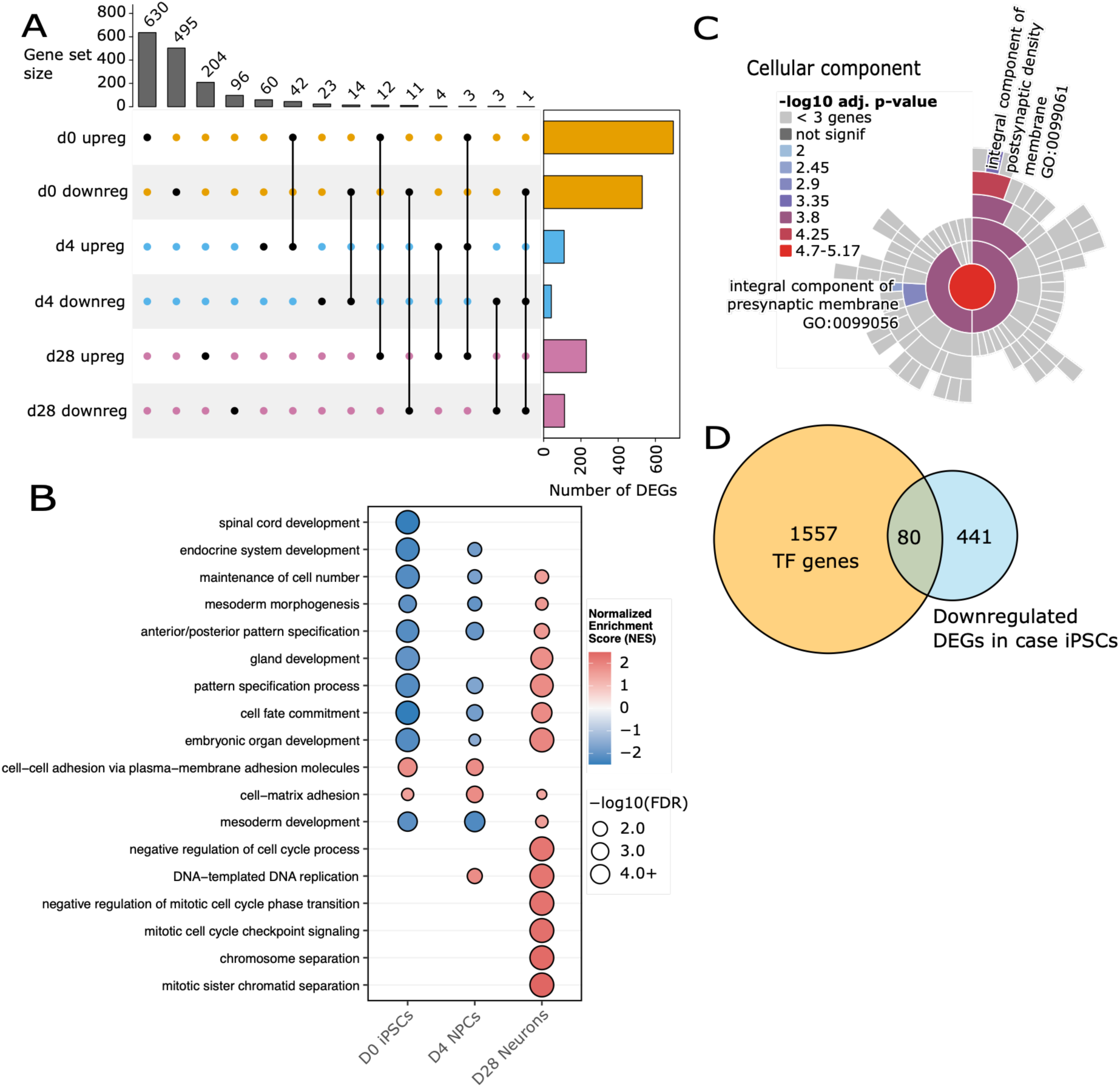
*INTS6* case iPSCs show downregulated developmental and transcription factor gene expression and neurons show downregulation of synaptic genes. (A) UpSet plot visualising the shared and unique sets of DEGs that are upregulated or downregulated at each timepoint. The vertical bars represent the intersection size, indicating the number of common DEGs between specified differentiation timepoints. The horizontal bars on the right show the total number of upregulated or downregulated DEGs for each timepoint. Notably, the largest intersections are observed between upregulated in iPSCs and upregulated in NPCs, and between downregulated in iPSCs and downregulated in NPCs. **(B)** Dotplot illustrating Biological Process terms from GSEA across iPSCs (D0), NPCs (D4), and neurons (D28). Terms were ranked by adjusted p-value within each timepoint (top 10 retained), and semantically redundant terms were collapsed using Wang similarity (GOSemSim, cutoff = 0.7). Dot colour denotes normalized enrichment score (NES); dot size denotes significance (−log10 FDR). Blank cells indicate non-significant enrichment (FDR > 0.05). The term with the highest significance was selected as the representative term for each cluster to facilitate biological interpretation. **(C)** Sunburst plots of enriched cellular component GO-terms from significantly downregulated DEGs in *INTS6* case D28 neurons. Plot made using SynGO v1.3. **(D)** The DEGs that are significantly downregulated in cases are enriched for transcription factors according to a curated list of known human transcription factors (OR = 2.44, p = 6.349 × 10^−11^).

To analyse which biological processes were affected by the DEGs between cases and controls across differentiation, we performed gene set enrichment analysis (GSEA). In case iPSCs, there was reduced expression of genes related to developmental and differentiation-associated biological processes (p < 0.05, NES < -1.5) including cell fate commitment (GO:0045165), spinal cord development (GO:0021510) and embryonic organ development (GO:0048568), alongside depletion of stem cell-related processes **(Table S7, Fig. 3B).** In case NPCs, downregulated genes were enriched for developmental pathways (p<0.05, NES < -1.5), including mesoderm development (GO:0007498), pattern specification process (GO:0007389) and cell fate commitment (GO:0045165), whereas upregulated genes were enriched for cell adhesion pathways, including cell–matrix adhesion (GO:0007160) and cell– cell adhesion via plasma membrane adhesion molecules (GO:0098742) **(Table S8).** In case neurons, cell cycle-associated pathways showed upregulation (p<0.05, NES > 1.5), including regulation of mitotic sister chromatid separation (GO:0010965), chromosome separation (GO:0051304), and DNA-templated DNA replication (GO:0006261). Neuronal signalling pathways such as chemical synaptic transmission (GO:0007268) were downregulated **(Table S9, Fig. 3B).** This suggests that partial LoF of *INTS6* can disrupt the developmental trajectory of neuronal cells, characterised by persistent impairment of cell fate specification and developmental programs in iPSCs and NPCs, transient alterations in cell adhesion in NPCs, and a shift towards activation of cell-cycle and proliferation-related pathways in neurons.

Guided by the GO-term enrichment for synapse, we used SynGO^44^ to determine whether any specific synaptic functions were affected by the *INTS6* variant. We found that significantly downregulated genes in neurons were enriched for synaptic terms such as process in the synapse (SYNGO:synprocess, p.adj = 6.7×10^−5^, 21 genes/1,252 background genes), postsynaptic density membrane (GO:CC:0098839, p.adj = 1.0×10^−4^, 7 genes/145 background genes), and regulation of presynaptic cytosolic calcium levels (SYNGO:presynprocess:GO:0099509, p.adj = 1.8×10^−4^, 4 genes/29 background genes) **(Fig. 3C, Fig. S3I, Table S10).** Downregulation of synaptic and postsynaptic functions in case neurons suggests that the transcriptional effects of partial loss of *INTS6* could converge on the synapse, which is highly consistent with genes and mechanisms previously proposed in neurodevelopmental and neuropsychiatric disorders ^45–48^, particularly affecting excitatory glutamatergic neurons^48,49^.

Given the role of *INTS6* in modulating transcription, we separately assessed the overlap between genes significantly upregulated or downregulated in cases with a curated list of all known human transcription factors (TFs)^51^. We found an enrichment of TFs in downregulated DEGs in case iPSCs (OR=2.44, 95% CI: 1.89-3.12, p=6.349 × 10^−11^), but not in NPCs or neurons **(Fig. 3D, Fig. S3F-H)**. This suggests that the *INTS6* partial LoF could exert its effects via impairments in transcriptional regulation at early developmental stages in the transcriptionally active iPSCs. Still, some of the downregulated TFs (n=80/1,637) from D0 remained downregulated at D4 and D28 without reaching statistical significance, suggesting a gradual attenuation of this transcriptional signature **(Fig. S3G-H)**.

Previously, *Ints6* LoF in rodents was found to disrupt RNAPII regulation, leading to excessive RNAPII activity at key cell cycle regulators, causing upregulation of cell cycle pathways, leading to impaired neurogenesis. To evaluate the conservation of transcriptional changes between our human iPSC-based *INTS6* models and mouse models, we compared our DEGs (p < 0.05) with those from *Ints6* nervous system conditional knockout models profiled by Peng *et al.* (2025)^22^ at different development stages: E15.5 *Ints6* nervous system conditional knockout (cKO) and 2-month conditional heterozygous knockout (cHET). Of the 30,636 genes with detectable transcripts in the cKO dataset, 16,483 (53.8%) mapped to human orthologs, and of the 31,668 genes in the cHET dataset, 16,156 (51.0%) mapped to human orthologs. Of the 2,900 mouse cKO DEGs, 2,442 (84.2%) and of the 826 mouse cHET DEGs, 732 (88.6%) mapped to human orthologs. Comparing shared orthologous DEGs, there were 144 shared DEGs between cKO and D0 (2,900 cKO DEGs, 1,208 D0 DEGs; hypergeometric test, BH-adjusted p = 0.89), 15 shared DEGs between cKO and D4 (150 D4 DEGs; p = 0.88), and 46 shared DEGs between cKO and D28 (323 D28 DEGs; p = 0.88). Between cHET (826 DEGs) and D0, there were 59 shared DEGs (p = 0.30); 4 shared DEGs between cHET and D4 (p = 0.88); and 13 shared DEGs between cHET and D28 (p = 0.88) **(Table S11, Fig. S3J).** None of the six comparisons showed a significant excess of shared DEGs beyond what would be expected by chance after correction for multiple testing. Direction concordance among shared DEGs ranged from 32.6% (cKO vs D28) to 76.9% (cHET vs D28); however, none of the six comparisons showed significant concordance after correction for multiple testing (binomial test, BH-adjusted p > 0.05 for all comparisons) **(Table S11).** Despite the lack of significant DEG concordance, there was a significant positive correlation of the log_2_FC values of the 13 shared DEGs between cHET and D28 (Pearson R = 0.67, p = 0.013), but not for the other comparisons **(Fig. S3K).**

Among the concordant genes in the cHET vs D28 comparison, several are linked to synaptic and neuronal functions. *ITPR1* (upregulated in both), which is involved in calcium signalling in neurons, *CPEB3* (downregulated in both) that regulates local protein synthesis in synapses, *SIPA1L1* (upregulated in both) that regulates dendritic spine and synaptic structure organisation, and protocadherins *PCDHGA3* and *PCDHGC4* (downregulated in both), which regulate dendritic arborization and synapse formation in the cortex. **S**ome genes followed opposite regulation trends, as is the case of *OLFM* (downregulated in NFID, upregulated in Peng *et al.*), which regulates synapse formation and neural development **(Table S11).** Overall, the overlap of these genes suggests relevant effects of *INTS6* LoF on synaptic function and neuronal connectivity, in both mouse and human NDD models. Together, these findings suggest both similarities and potential species divergence in the effects of *INTS6* LoF.

### The *INTS6* variant associates with increased intron retention in cases

Given the Integrator’s role in transcriptional modulation through snRNA 3’-end processing^50^ and the involvement of pathogenic variants in components of the major and minor spliceosomes and associated endonucleases in human disease ^17,51–55^, including NDDs^12,13^, we next asked if the *INTS6* variant affects splicing. In the alternative splicing analysis, splice clusters were defined as groups of overlapping introns sharing a common donor or acceptor site, reflecting splicing at a shared genomic location. Analysis was restricted to junctions supported by at least 50 reads in iPSC and NPC samples and at least 20 reads in neuron samples (which included rat astrocytes and correspondingly smaller human-aligned libraries). Differential splicing analysis revealed significant alternative splicing events across all timepoints in cases (p.adj < 0.05), affecting 357 splice clusters and 384 genes in iPSCs, 423 clusters and 474 genes in NPCs, and 149 clusters and 178 genes in neurons (**Table S12**). Among the splicing events overrepresented in cases, junctions with known donor and acceptor sites were the most abundant in iPSCs (77.1%), NPCs (74.9%) and neurons (67.5%). Similarly, junctions with known donor and acceptor sites were the most abundant among the splicing events underrepresented in cases: iPSCs (76.2%), NPCs (72.1%) and neurons (63.2%) **(Table S12).** Together, these findings indicate that known splice junctions predominate among both over- and underrepresented splice events, with their relative abundance decreasing from iPSCs to neurons.

To further investigate the effects of the *INTS6* partial LoF on RNA processing, we studied differences in intron retention (IR) between case and control iPSCs by analysing 17,304 exons in 5,323 genes that met sufficient read coverage and IRFinder quality control criteria (see Methods). IR was significantly altered in 1,101 exons across 720 genes. Notably, all but one gene (719) had significantly increased IR in cases relative to controls. Almost all tested exons (N = 1,100/1,101, p = 4.42 × 10^−31^, OR = 80.2) had more intronic reads in cases than controls **(Fig. 4A, Table S13).** A transcriptome-wide splicing defect in iPSCs was further supported by the associated p-value distribution that deviated from the null expectation (ΔPSI> 5%, p.adj <0.05) **(Fig. 4B).** Among the 719 genes with significant IR in cases, we identified the histone lysine demethylase *KDM5B,* a causative gene for autosomal NDDs such as ID, ASD, and motor speech disorders^56^ **(Fig. 4C, Fig. S4A)**. In case NPCs and case neurons, IR was not significantly higher than in controls after multiple testing **(Fig. 4B)**.

**Figure 4:**
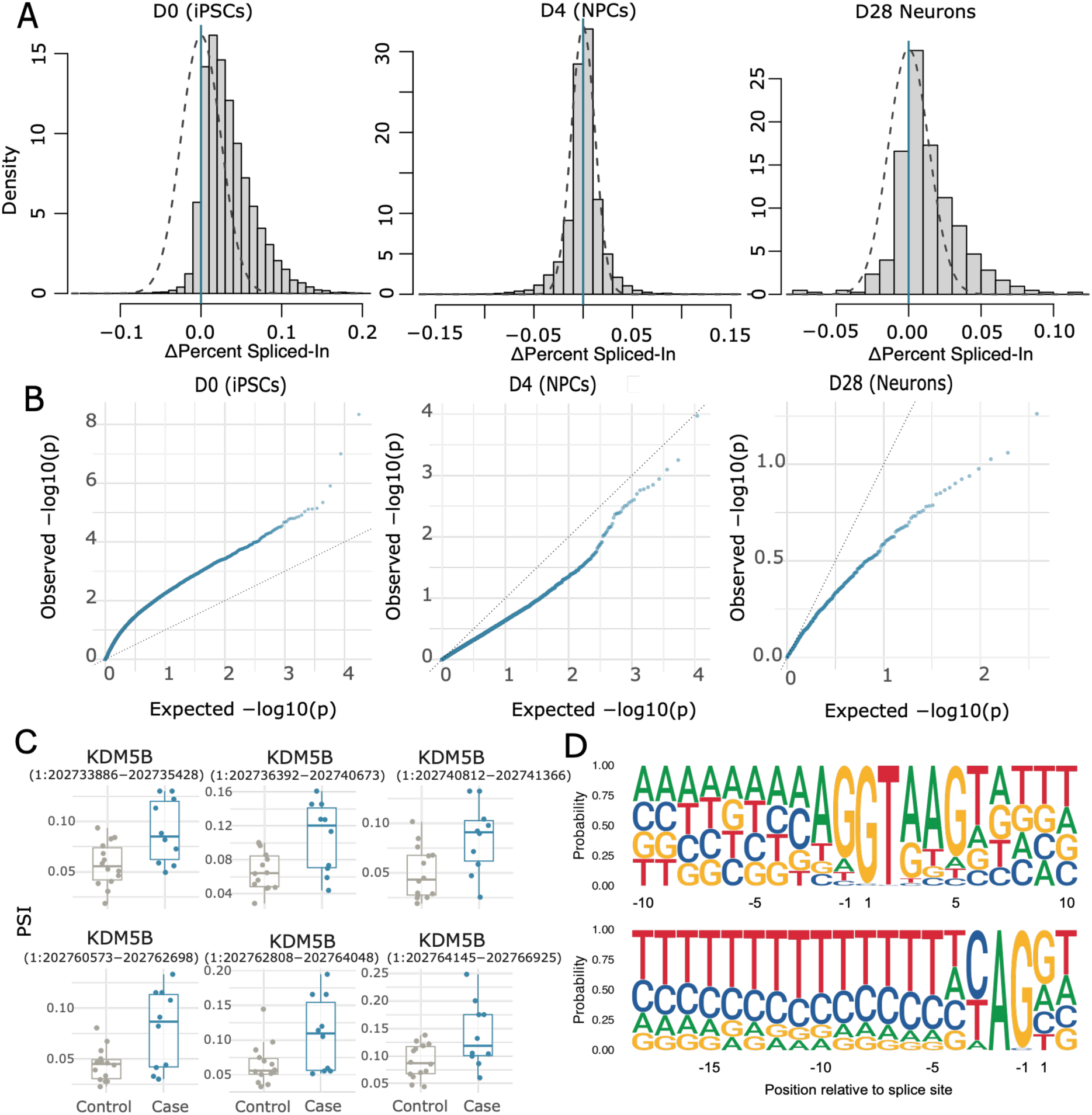
The *INTS6* heterozygous variant is associated with intron retention. **(A)** Histograms showing that distributions of intron retention scores are shifted toward higher values in cases compared to controls in iPSCs (p < 2.2 × 10⁻¹⁶, Wilcoxon test), NPCs (p = 8.34 × 10⁻¹¹) and neurons (p < 2.2 × 10⁻¹⁶). **(B)** Q-Q plots show an excess of small p-values relative to the null expectation for differential intron retention in cases versus controls in iPSCs, but not in NPCs or neurons. **(C)** Boxplots showing that *KMD5B* (an NDD gene) has six introns that are significantly more retained in cases than in controls (p < 0.05). **(D)** The splice sites of introns with significant intron retention (N=1,100) show normal U2-type nucleotide distribution both at 5’ (upper) and 3’ (lower) splice sites.

However, as observed in iPSCs, the distribution of ΔPSI values indicated a bias towards more retained introns in cases in NPCs and neurons (Wilcoxon test, p < 2.2 × 10^−16^) **(Fig. 4A).** The splice sites of the introns with significantly increased IR in cases (N = 1,100) were canonical U2-type splice sites **(Fig. 4D)**, further suggesting that IR arises independently of splice site mutations. Although we expected IR genes to be longer and hence more susceptible to splicing defects, we found that gene length was weakly negatively correlated with ΔPSI (Pearson R = -0.031, p = 0.0253) **(Fig. S4B).**

To understand if impaired splicing was associated with differential gene expression, we analysed the overlap between IR genes and DEGs in case iPSCs. Although no direct overlap was observed **(Table S4)**, both gene sets converged at the level of GO-enrichments, particularly in biological process and cellular component terms. GO-term enrichment revealed that genes with significant IR in case iPSCs (N=720) were enriched (p < 0.05, fold changes > 1.5) for translation (GO:0006412) and cytoplasmic translation (GO:0002181), indicating increased translational activity. Genes with IR also had enrichment of DNA metabolic and repair pathways, including DNA metabolic process (GO:0006259), and DNA repair (GO:0006281). Strong enrichment was also observed for genome stability and proliferative networks, including cell cycle (GO:0007049), mitotic cell cycle process (GO:1903047), chromosome segregation (GO:0007059) and chromatin organization (GO:0006325). The genes with significant IR were also involved processes relevant to the Integrator complex including RNA 3’-end processing (GO:0031123), RNA splicing (GO:0008380), regulation of transcription elongation by RNA polymerase II (GO:0034243), RNA localization (GO:0006403) and ribonucleoprotein complex (GO:1990904) **(TableS14).** Together, these findings suggest that while increased translation may reflect the high proliferative state of iPSCs, the enrichment of chromatin organisation, cell cycle and RNA processing pathways points to genome dysregulation relevant to NDDs^47^.

### Engineered knock-in iPSCs recapitulate the transcriptomic signatures associated with the heterozygous *INTS6* variant

To determine whether the observed transcriptomic changes were causally linked to the *INTS6* variant, we used CRISPR-Cas9 genome editing to introduce the variant into an unaffected family control iPSC line and to correct the variant in a family carrier iPSC line **(Fig. 5A).** For the knock-in, we collected both heterozygous (KI, N=3 clones) and homozygous clones (KO, N=1 clone) **(Table S15)**. Immunocytochemistry of the engineered iPSC KI and KO lines showed expression of markers of the undifferentiated state (OCT4, Nanog and Tra-1-81) **(Fig. S5A).** In contrast, immunocytochemistry of D4 NPCs for the same lines showed expression of Nestin, PAX6 and SOX1, and absence of OCT4 **(Fig. S5B)**. This marker profile indicates a normal loss of pluripotency and progression toward a neurogenic state characterised by expression of neuron progenitor features, even in the CRISPR KO. This differentiation marker profile suggests that the *INTS6* LoF does not affect *Ngn2* NPC differentiation potential, although heterozygous or homozygous loss of *INTS6* could still affect synaptic processes in neurons.

**Figure 5:**
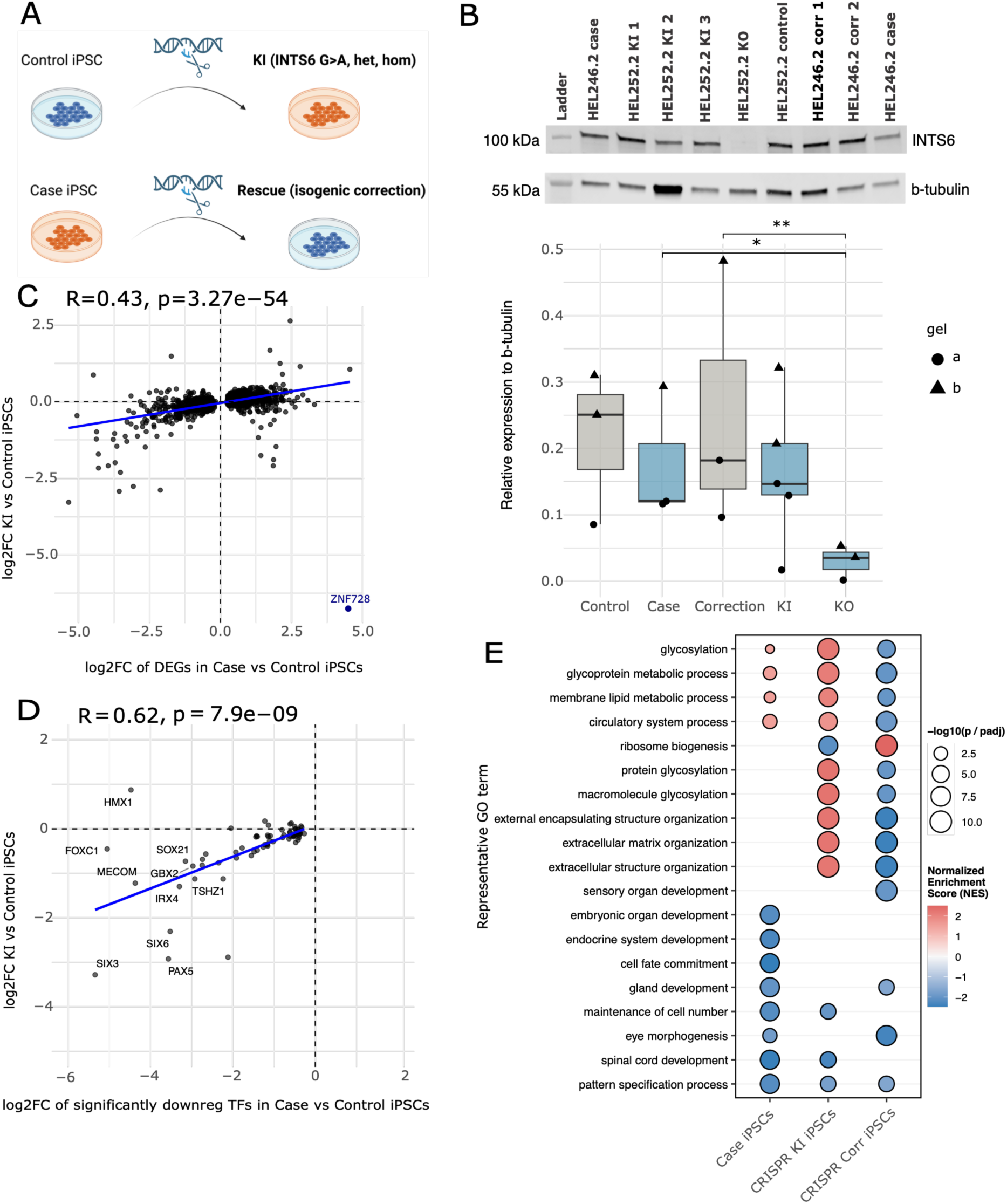
CRISPR-Cas9 engineered iPSCs validate the protein and transcriptional effects of *INTS6* LoF. **(A)** Schematic of CRISPR-Cas9 genome editing depicting correction of the *INTS6* chr13:51378376 G>A variant in a patient-derived iPSC line and heterozygous/homozygous knock-in of the same variant into a control iPSC line. **(B)** Western blot showing reduced expression of the 100 kDa INTS6 protein in the CRISPR KO iPSC line, with a representative blot (upper panel) and quantified band intensities normalized to β-tubulin across all replicates (lower panel). Boxplots show median and interquartile range (IQR), with whiskers extending to the most extreme value within 1.5 × IQR; points represent biological replicates, with point shape indicating gel (technical replicate). Statistical significance was assessed by linear mixed-effects model with Tukey-adjusted post-hoc comparisons (*p < 0.01). **(C)** Scatterplot showing modest but significant correlation (Pearson R = 0.43, p = 3.27×10^−54^) between log_2_FC of DEGs in case vs control iPSCs and log_2_FC of the same genes in CRISPR KI vs control iPSC**. (D)** The set of 80 downregulated TF genes in case iPSCs are also downregulated in the CRISPR KI iPSCs (Pearson R = 0.62, p = 7.9×10^− 9^). Highlighted genes include *SIX3, SIX6* and *FOXC1*, which contribute to neural progenitor proliferation. **(E)** Dotplot illustrating Biological Process terms from GSEA across case, CRISPR KI and CRISPR corrected iPSCs. Terms were ranked by adjusted p-value within each timepoint (top 10 retained), and semantically redundant terms were collapsed using Wang similarity (GOSemSim, cutoff = 0.7). Dot colour denotes normalized enrichment score (NES); dot size denotes significance (−log10 FDR). Blank cells indicate non-significant enrichment (FDR > 0.05). The term with the highest significance was selected as the representative term for each cluster to facilitate biological interpretation.

Western blot analysis revealed a significant overall effect of genotype on INTS6 protein levels (F(4,11) = 6.05, p = 0.0079, linear mixed-effects model with gel as a random effect). Post-hoc Tukey-adjusted pairwise comparisons showed a significant decrease in INTS6 protein in KO compared to case (N_lines_ = 3) (estimate = 0.269, t(11) = 4.22, adj. p = 0.0100) and correction (N_clones_ = 2) (estimate = 0.276, t(11) = 4.33, adj. p = 0.0084) lines, with a marginal reduction compared to KI (N_clones_ = 3) (estimate = 0.182, t(11) = 3.20, adj. p = 0.0525) and a suggestive reduction compared to control (N_lines_ = 3) (estimate = 0.179, t(11) = 2.88, adj. p = 0.0890) **(Fig. 5B).** Similarly to the case lines, the KI appeared to differ only modestly in INTS6 protein levels from case, correction, or control lines (all adj. p > 0.47). As seen earlier in the heterozygous cases **(Fig. S1C)** and distinct from the overexpressing cells, no truncated protein at 54.6 kDa was detected even in the KO line **(Fig. S5C).**

We performed RNA-sequencing on the CRISPR-edited iPSCs to assess the impact of genetic modifications on gene expression. The PCA showed that the CRISPR lines, including isogenic corrected clones (N=2), KI clones (N=3), and a KO clone (N=1), clustered with cases and controls, indicating that the CRISPR editing process did not introduce major additional transcriptional effects beyond the expected parental background **(Fig. S5D).**

Along PC1 (34.8% variance explained), the CRISPR lines clustered with their respective parental iPSC line and not with their genotype **(Fig. S5E),** in line with the observed subtle transcriptomic influence of the *INTS6* variant in case lines. In line with results from case iPSCs, the CRISPR KO of *INTS6* had little effect on *INTS6* expression, with no detectable differences in mRNA expression between cases, controls, CRISPR-corrections, KI or KO iPSCs **(Fig. S5F).** In the CRISPR KO iPSC line, there is a reduced expression of *INTS6* exons after the predicted stop-codon, consistent with the LoF mechanism **(Fig. S5G).** To test whether the KI lines recapitulated transcriptional signatures observed in cases, we considered the 1,208 genes that were differentially expressed in the iPSCs of the original case vs control dataset (adj. p < 0.05). Effect sizes showed a modest but significant correlation between the two groups (Pearson R = 0.43, p = 3.27×10^−54^) **(Fig. 5C, Table S16).**

Unlike the iPSC case vs control comparison, CRISPR KI and KO iPSCs did not recapitulate the IR phenotype seen in cases **(Table S17)**. Still, the downregulation of the 80 TF genes in case iPSCs **(Fig. 3D)** was also visible between in KI iPSCs (Pearson R = 0.62, p = 7.9 × 10⁻⁹) **(Fig. 5D).** Among the TFs, we identified *SIX3*, *SIX6* and *FOXC1*, which regulate neural progenitor proliferation during neurodevelopment^57–59^. As in case iPSCs, the GSEA biological process maintenance of cell number (GO:0098727) was downregulated in CRISPR KI iPSCs but returned to baseline in the CRISPR corrected iPSCs **(Table S18, Fig. 5E).** This suggests that the *INTS6* variant may affect cell proliferation, consistent with disrupted proliferation-differentiation balance in NDDs^60^.

### *INTS6* LoF does not lead to measurable changes in major spliceosome U-snRNA expression levels or RNA polymerase II phosphorylation

Previous literature indicates that the phosphatase and endonuclease modules of the Integrator have distinct roles in transcription. The phosphatase module is involved in modulating transcription through RNAPII dephosphorylation, while the endonuclease module is responsible for snRNA 3’-end processing^39,40^. We investigated whether the *INTS6* variant affects RNAPII phosphorylation in iPSC as a potential contributor to the transcriptional dysregulation observed in cases. We hypothesized that loss of one *INTS6* copy could disrupt the phosphatase module function resulting in hyperphosphorylation of RNAPII-CTD serine residues, such as Ser2 **(Fig. 6A).** However, Western blot analysis showed no significant overall effect of genotype on RNAPII-CTD pSer2 protein levels (F(4,16) = 1.55, p = 0.24, linear mixed-effects model with gel as a random effect). Pairwise comparisons between groups did not reach statistical significance (all adj. p > 0.34, post-hoc Tukey), including between case and control (estimate = 0.306, t(14.4) = 1.94, adj. p = 0.34), case and correction (estimate = −0.042, t(13.5) = 0.19, adj. p > 0.99), case and KI (estimate = 0.103, t(13.5) = 0.48, adj. p = 0.99), and case and KO (estimate = −0.073, t(13.5) = 0.34, adj. p = 0.99) **(Fig. 5B).**

**Figure 6:**
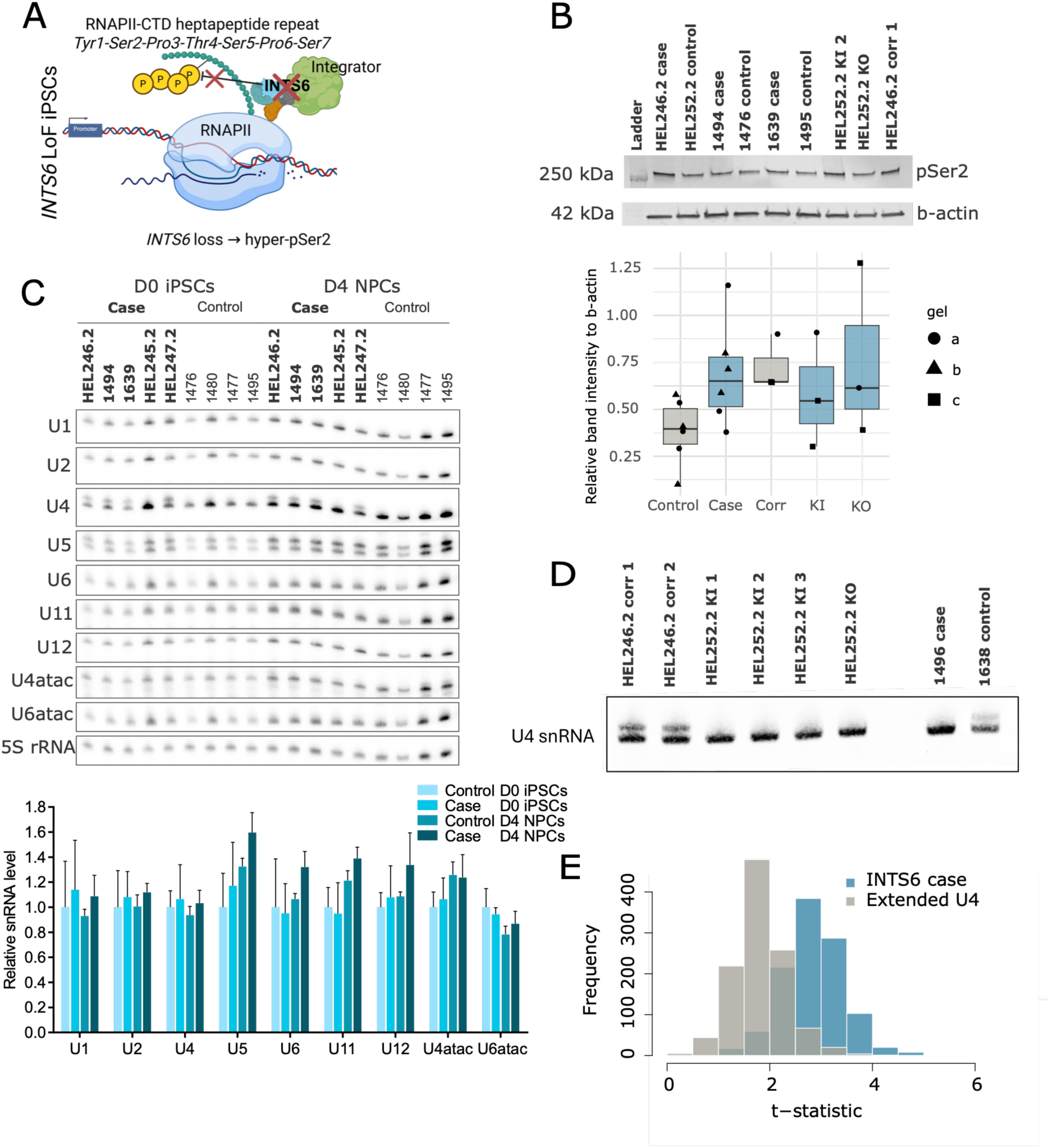
The *INTS6* variant does not significantly affect major spliceosome U-snRNAs or RNAPII phosphorylation. **(A)** Schematic showing the hypothetical effect of loss of Integrator phosphatase module function via *INTS6* LoF on hyperphosphorylation of the Ser2 residue on RNAPII-CTD. Panel A was created using BioRender. **(B)** Western blot showing pSer2 RNAPII-CTD levels (upper panel) and quantified band intensities normalized to β-actin (lower panel). Boxplots show median and interquartile range (IQR), with whiskers extending to the most extreme value within 1.5 × IQR; points represent biological replicates, with point shape indicating gel (technical replicate). No significant differences in Ser2 phosphorylation were observed between groups (linear mixed-effects model with Tukey-adjusted post-hoc comparisons; all adj. p > 0.05). **(C)** Northern blot showing major and minor spliceosome U-snRNA levels in cases and controls (upper panel) and quantifications relative to ribosomal 5S rRNA (lower panel) showing no significant differences in U-snRNA levels between cases and controls within D0 iPSCs and D4 NPCs. **(D)** Northern blot of U4-snRNA in iPSCs showing that the isogenic corrected clones do not remove the U4 extended form (left) and that the heterozygous knock-in (KI) or homozygous knock-in (KO) of the *INTS6* variant does not produce the U4 extended form (right). **(E)** Histogram showing frequency of ΔPSI IR scores (IRFinder) shift to higher values in *INTS6* case iPSC compared to iPSC lines with the extended form of U4, suggesting that IR is more likely to be a consequence of the *INTS6* mutation rather than the extended form of U4-snRNA (p = 9.88×10^−289^, Welch two-sample t-test).

Given the observed IR in *INTS6* cases, we examined the potential effects on major and minor spliceosome U-snRNAs using Northern blotting. We quantified relative RNA expression levels of U1, U2, U4, U5, U6, U11, U12, U4atac, U6atac, as well as 7SK snRNA and P-TEFb in D0 iPSCs and D4 NPCs for cases (N=5) and controls (N=4). Ribosomal 5S rRNA was used for normalization (probes detailed in **Table S19).** None of the pairwise comparisons between cases and controls for each snRNA were statistically significant (p > 0.05) **(Fig. 6C, Table S20).** While the *INTS6* variant showed no significant effect on overall U-snRNA levels between cases and controls at any timepoint, Northern blot analysis revealed an extended form of U4-snRNA in 4 out of 5 case iPSC and NPC lines tested **(Fig. 6C).** Initially, we thought the extended U4-snRNA may indicate an unprocessed form with a longer 3’-tail as a downstream consequence of the *INTS6* variant. We tested this hypothesis using the CRISPR-Cas9 edited iPSC lines and found that correcting the *INTS6* variant did not rescue the extended U4, while KI or KO lines did not present the extended U4.

Additionally, the *INTS6* case iPSC line 1469 did not show the extended U4, while the control line 1638 presented it **(Fig. 6D),** further suggesting that the extended U4 was unrelated to the *INTS6* variant.

Considering the presence of the extended U4-snRNA in 4 out of 6 case cell lines, and the significantly higher IR in case iPSCs compared to control iPSCs, we assessed whether IR frequency was more elevated in iPSC lines with the *INTS6* variant or in those exhibiting the extended U4 form. We found that IR frequency was significantly higher in *INTS6* case iPSCs compared to iPSCs with the extended form of U4, suggesting that the observed IR is more likely to be a consequence of the *INTS6* mutation rather than the extended form of U4-snRNA (p = 9.88×10^−289^, Welch two-sample t-test). **(Fig. 6E).**

## Discussion

Disruption of Integrator complex phosphatase activity has been linked to diseases including NDDs, but how the subunit *INTS6* is implicated in these diseases in humans remains poorly understood. Here, we report a novel *INTS6* c.1465C>T p.Arg489* variant that causes predominantly mild ID in a Finnish family with six affected individuals **(Fig. 1).** Using human iPSC-derived neuronal models, we demonstrate that *INTS6* haploinsufficiency leads to genome-wide dysregulation of RNA processing, including increased intron retention at canonical splice sites. The genes presenting significant intron retention in cases are enriched for the biological processes of cell cycle, RNA-3’-end processing, RNA splicing, and regulation of transcription elongation by RNA polymerase II **(Fig. 4)**. We found that in case iPSCs and NPCs, the *INTS6* partial LoF caused transcriptional changes that were associated with impaired developmental and cell fate specification programs. Later in differentiation, mature neurons in cases presented aberrant activation of cell cycle-related pathways and reduced neuronal signalling signatures. We also found downregulation of transcription factor genes in early development in case iPSCs, and downregulation of pre- and synaptic functions in case neurons **(Fig. 3)**. Engineered mutations partially recapitulated these findings, with KI lines mirroring the transcriptomic profile of cases and suppression of biological processes related to maintenance of cell number, reinforcing the role of *INTS6* in development and neuron function **(Fig. 5)**.

The Integrator complex has been implicated in NDDs, with reported germline pathogenic variants in *INTS1*^16,61^*, INTS8*^18^*, INTS11*^19^, and *INTS13*^20^. *INTS10* depletion has been shown to shift cellular identity toward mesenchymal states^62^, while *INTS8* loss enhances nascent transcription and accelerates neuronal gene expression, ultimately compromising the neural progenitor pool^18^. Recently, 21 *de novo* monoallelic variants in *INTS6* have been linked to NDDs, with most common clinical features being speech and language impairment and ID. These features were also observed in the *INTS6* c.1465C>T p.Arg489* cases described here, which likewise showed a male predominance^22^. Pathogenic variants in *INTS6* are also associated with clinical variability, a feature typically observed in NDDs. In a non-human model, Peng *et al.* (2025) found that *Ints6* deficiency in mice disrupts PP2A-RNAPII regulation, leading to excessive RNAPII activity, resulting in activation of cell cycle pathways, which we also saw in *INTS6* partial LoF neurons. The partial *INTS6* LoF did not alter the capacity to generate excitatory neuron differentiation, which is partly consistent with previous findings in heterozygous *Ints6* nervous system cKO mouse data^22^. This is consistent with known NDD mechanisms of synaptic function, neuronal connectivity and activity-dependent plasticity^45,63,64^. Overall, our findings suggest that future experiments should investigate the effects of *INTS6* LoF on synaptic biology.

We used *Ngn2* overexpression-driven differentiation to generate glutamatergic neurons, thereby modelling excitatory neuronal identity and intermediate progenitor populations. Transcriptomic analysis indicated minimal batch effects and high reproducibility, as supported by strong correlations between technical replicates and clear case-control clustering across timepoints in the RNA-seq data. However, *INTS6* may have broader effects across other neuronal lineages and developmental stages, necessitating future studies that incorporate alternative differentiation models to capture the full spectrum of *INTS6* function and compensatory mechanisms in NDDs.

Notably, the absence of detectable differences in INTS6 protein levels or truncated protein by Western blot may reflect compensatory mechanisms due to heterozygosity. The CRISPR KO iPSC line produced no detectable full-length or truncated INTS6 protein, whereas HEK293T cells overexpressing the mutant INTS6 allele produced no full-length protein but did yield a truncated ∼55 kDa product, confirming that the mutant transcript is translated.

Since *INTS6* mRNA levels were unaffected in both CRISPR KI and KO cells, this indicates that the mutant transcript largely escapes nonsense-mediated decay, and that protein loss instead reflects post-translational instability of the truncated product, detectable only when protein production is forced beyond normal degradation capacity, as with HEK293T overexpression. This raises questions about the threshold of *INTS6* expression necessary to observe phenotypic effects, particularly in the context of transcriptional regulation and synaptic biology. Future studies should explore whether this compensatory effect can mask the functional consequences of *INTS6* haploinsufficiency, especially in the context of mild ID, and how it might influence the observed dysregulation in RNA processing and synaptic gene expression.

Defective Integrator activity has been shown to shape the transcriptome in human diseases through reduced pre-mRNA splicing efficiency and disrupted gene expression of RNAPII-associated genes^53^. While *INTS6* cases had significantly higher intron retention globally, specifically in genes enriched for RNA processing, it was less pronounced in neurons, possibly reflecting reduced transcriptional activity in post-mitotic neurons. Notably, dynamic IR is common during cell differentiation and may reflect regulated transcript processing^65^.

Moreover, in our CRISPR-Cas9 edited lines, IR was not significantly higher in heterozygous KI or KO iPSCs compared to CRISPR corrections. This may be partially attributed to the limited number of parental lines used in CRISPR editing, potentially yielding insufficient power to detect IR differences between conditions even when using multiple edited clones for one iPSC line. Additionally, patient genetic background may contribute to the molecular phenotypes observed here beyond the variant of interest. Our results indicate that *INTS6* LoF does not alter major or minor spliceosome U-snRNA levels, in line with independent functions of the Integrator phosphatase and endonuclease modules in transcription regulation^39,40,66^. In this context, the fate of IR-containing transcripts remains to be explored.

Transcription pause-release is a critical process in timing and controlling gene expression. The Integrator phosphatase module is involved in suppressing transcription of coding genes^40^. Previous research suggests that depleting *Ints6*^25^ or *Ints8*^27^ in *D. melanogaster* resulted in global upregulation of gene expression, aligning with the role of the phosphatase module in regulating RNAPII pausing and transcriptional output. Related to *INTS6* partial LoF, we also detected a consistent and significant upregulation bias in gene expression.

Linked to this, the phosphatase module is involved in dephosphorylating RNAPII, and if defective, can lead to hyperphosphorylation of serine residues (Ser2, Ser5, Ser7) on RNAPII-CTD, which typically occurs during transcription elongation^25,27,39^. However, we did not detect significant Ser2 hyperphosphorylation in cases or CRISPR-Cas9 KI or KO iPSCs compared to control and isogenic correction iPSCs measured by Western blot. As Western blotting has limited sensitivity for detecting subtle quantitative differences^67^, particularly in low-abundance proteins, we cannot exclude modest changes in Ser2 phosphorylation. This suggests that compensatory mechanisms or technical limitations may obscure the expected RNAPII phosphorylation alterations in our experimental models.

Whether the Integrator complex regulates transcription globally or primarily acts on specific gene targets in humans remains an open question. Recent evidence suggests that the phosphatase module is essential for transcribing a select group of genes, with protein phosphatase 2A recruitment fine-tuning transcription termination efficiency^25^. In *D. melanogaster* cells, *Ints6* overexpression upregulated genes enriched in axon generation, protein folding and heat response pathways^25^. Conversely, *Ints8* depletion upregulated immediate-early response genes^18^, many of which encode TFs. In our human models, however, TFs and genes involved in cellular differentiation were significantly enriched among downregulated genes in *INTS6* case iPSCs. Moreover, *INTS6* regulated distinct sets of genes at each developmental timepoint, suggesting transcriptional effects are context dependent. This highlights the complexity of the role of *INTS6* in transcriptional regulation in different species and cell types.

In summary, this study links a *INTS6* partial LoF variant with transcriptional dysregulation in cellular models of mild ID. Using human iPSC-derived neurons, we found that the heterozygous *INTS6* variant c.1465C>T, p.Arg489* leads to significant transcriptome-wide intron retention in genes enriched for transcription regulation, and downregulation of genes involved in transcription, cell fate specification and neuronal function. Although defective *INTS6* is likely to impair transcriptional regulation, compensatory mechanisms may obscure its effects on RNAPII activity, particularly in individuals with mild ID. Together, these findings identify *INTS6* LoF as a contributor to mild intellectual disability and highlight the need for future studies to define the gene networks and RNAPII-dependent regulatory mechanisms governed by *INTS6* in humans. Understanding these molecular interactions will be essential for understanding the molecular basis of NDDs.

## Methods

### Samples

The NFID study (Northern Finland Intellectual Disability study) was approved by the ethical committees of Northern Ostrobothnia Hospital District and the Hospital district of Helsinki and Uusimaa. All participants and/or their legal guardians have provided a voluntary written informed consent to participate in the study and for iPSC research. All the cell lines are pseudonymized, and all donor and sequence data generated from the cell lines were handled in a secure and protected environment (IT Center for Science Ltd. - CSC ePouta) according to data protection regulations in the EU and Finland.

The cases have all been evaluated and examined clinically by multi-professional teams (cases described in **Table S1**). Standardized IQ tests that were used included different versions of the following tests: Wechsler Preschool and Primary Scale of Intelligence (WPPSI), Wechsler Intelligence Scale for Children (WISC) and Wechsler Adult Intelligence Scale (WAIS) for adults.

### iPSC reprogramming and characterisation

All iPSC lines were reprogrammed from peripheral blood mononuclear cells (PBMCs) with Sendai virus technology^68^, at the Harvard Stem Cell Institute or Biomedicum Stem Cell Center, University of Helsinki. The PBMCs were used to generate iPSCs by reprogramming with four recombinant Sendai viral vectors (CytoTune™, Life Technologies) expressing Oct4, Sox2, Klf4, and c-Myc. All iPSC lines were characterised with immunocytochemistry for markers of undifferentiated state *Nanog, Oct4* and *Tra181*. G-banded karyotypes were provided by Ambar Labs, Barcelona. These methods are detailed below, while iPSC metadata is described in **Table S2**.

### Human induced pluripotent stem cell culture

iPSCs were cultured on Matrigel (Corning, 354234) coated 35mm cell culture dishes (Thermo Scientific, 150460) in E8 basal medium (Gibco, A15169-01). Post-thawing, cell culture medium was supplemented with Rock-inhibitor (Selleck, Y-27632). iPSCs were stored in an incubator at conditions +37 °C, 5 % CO2. iPSCs were frozen in KnockOut serum replacement (KOSR, Thermo-Scientific, 10820-010) with 10% DMSO (Sigma, D2650).

### HEK293T cell culture

HEK293 cells were cultured on 100 mm cell culture dishes (Thermo Scientific, #130182) in DMEM High Glucose (Euro Clone, ECB7501L) containing 10% Foetal Bovine Serum (FBS) (Merck, #TMS-013-B), 1% penicillin-streptomycin (Lonza, #09-757F), and 1% L-Glutamine (Thermo Scientific, #25030149). Cells were passaged using 1x Trypsin (Thermo Scientific, #R001100) and stored in an incubator at conditions +37 °C, 5 % CO2.

### Rat primary astrocyte culture

Rat cortices were extracted from Wistar rat embryos at E17-18 under the permit for internal animal usage (KEK20-009) at the Neuroscience Center, University of Helsinki. After initial cell preparation, the purified cortical cells were plated on T75 flasks with 20 ml DMEM High Glucose (Euro Clone, ECB7501L) supplemented with 10 % fetal bovine serum (Gibco, 10500-064), 1 % L-Glutamine (BioWhittaker, BE17-605E) and 1 % Penicillin/Streptomycin (BioWhittaker, DE17-602E). After 7-8 days in culture with rat neurons reaching confluency, microglia and oligodendrocytes were removed by shaking the flask at 240 rpm for 6h. The cells were split using Trypsin (0.05%)-EDTA and media changed fortnightly.

### Neuronal differentiation and culture

Cortically patterned glutamatergic neurons were differentiated from iPSCs using *Ngn2* reprogramming and dual SMAD and WNT inhibition based on the Nehme et al. (2018) protocol^44^. iPSCs were infected overnight (24 h) with lentiviruses Tet-O-NGN2-PURO (Alstem, Cat CS-LV01, lot 220415, 3.65E+09 IFU/mL) and FudeltaGW-rtTA (Alstem, Cat CS-LV01, lot 220415, 2.39E+09 IFU/mL) using MOI=10. pTet-O-Ngn2-puro was a gift from Marius Wernig (Addgene plasmid # 52047 ; http://n2t.net/addgene:52047 ; RRID:Addgene_52047), and FUdeltaGW-rtTA was a gift from Konrad Hochedlinger (Addgene plasmid # 19780 ; http://n2t.net/addgene:19780 ; RRID:Addgene_19780).

Differentiation was induced with 2 μg/ml Doxycycline hyclate (Biogems, 2431450) in E8 medium (Gibco, A15169-01). Day 1 N2 medium consisted of DMEM/F-12 (Gibco, 21331-020), 1:100 Glutamax (Gibco, 35050-038), 1:100 N2 supplement (Gibco, 15410294, 1:67 20% Glucose), supplemented with 10 μM SB431542 (SB, Sigma, S4317), 100 nM LDN193189 (LDN, Sigma, SML0559), 2 μM XAV939 (Biogems, 2848932) and 2 mg/ml

Doxycycline. Day 2 N2 medium consisted of N2 medium supplemented with half of the day 1 supplements and 5 μg/ml puromycin (MP Biomedicals, 100552) to select for infected cells.

Day 3 media was the same composition as day 1 media. For final plating on day 4, the cells were detached with 1 mL Accutase (Gibco, 11599686) for 5 min and centrifuged at 300 rcf for 4 min before cell counting. Neurons were plated 1:1 with rat primary astrocytes onto cell culture dishes coated with poly-L-ornithine (Sigma-Aldrich) and 2x Matrigel using density of 60 000 cells/cm^2^. From day 4 onwards, the neurons were fed with Neurobasal medium (Gibco, 21103-049), 1:100 Glutamax, 1:200 MEM NEAA (Gibco, 11140-035), 1:67 20% Glucose, 1:50 B27 without vitamin A (Gibco, 12587001) supplemented with 10 ng/ml BDNF (PeproTech, 450-02), 10 ng/ml GDNF (PeproTech, 450-10) and 10 ng/ml CNTF (PeproTech, 450-13). Between days 7-8, proliferating cells were eliminated with NBM medium supplemented with 10 μM FUDR (Tocris, 4659). Half of the medium was changed every two days. Neurons were cultured for 28 days. The neuronal differentiations were carried out with support from the Neuroscience Center Neuronal iPSC core facility.

### Sample preparation

iPSCs, NPCs, and neurons were collected for downstream applications first by washing with 1 ml of RT DPBS to remove cell debris. 1 mL of +4 °C DPBS was added, and cells were detached carefully with a cell scraper (Thermo-Fisher Scientific) and transferred to 1.5 mL Eppendorf tubes and placed on ice. The cell suspensions were centrifuged at 1000 g for 7 minutes in +4 °C, supernatant aspirated, and pellets flash frozen on dry ice, and used for RNA extraction for bulk RNA-sequencing and Northern blotting, and protein extraction for Western blotting.

### Immunocytochemistry

D0 iPSCs, D4 Ngn2 neuron progenitor cells (NPCs), and D28 neurons co-cultured with rat primary astrocytes were fixed with 4% formaldehyde for 20 min and washed twice with DPBS. The cells were permeabilized with 0.1% Triton X-100 (Merck, T8787) in DPBS for 15 min at RT (except for cell surface marker Tra-1-81) followed by washing 3x with PBS-T (0.1% Tween-20 (Sigma, P1379) in DPBS. Non-specific binding sites were blocked using 5% BSA (Bovine Serum Albumin: BioWest, P6154) in PBS-T for 2 hours at RT. For D0 iPSCs, primary antibodies anti-Oct3-4 mouse (1:400, Santa Cruz, sc-5279), anti-Nanog rabbit (1:250, Cell Signaling Technology, 4903) and anti-TRA-1-81 (1:500, Cell Signaling Technology, MA1-024). For D4 NPCs, primary antibodies against Pax6 rabbit (1:400, Cell Signaling Technologies, 60433S), Oct3-4 mouse (1:400, Santa Cruz, sc-5279), Nestin rabbit (Abcam, ab105380) were used. For D28 neurons, primary antibodies against MAP2 chicken (1:500, Abcam, Ab92434), CUX1 mouse (1:500, Abcam, ab54583), VGLUT1 rabbit (1:300, Sigma, v0389-200), and GAD67 mouse (1:500, Abcam, ab26116) were used. Primary antibodies were diluted in 5% BSA in PBS-T and incubated overnight at +4°C. The cells were washed 3 x with DPBS. Secondary antibodies were donkey anti-rabbit AF488 IgG (H+L) (Invitrogen, A21206), donkey anti-mouse AF594 IgG (H+L) (Invitrogen, A21203), and goat anti-chicken Alexa Fluor 568 (Invitrogen, A11041). Secondary antibodies were diluted in 5% BSA in PBS-T and incubated for 1 h at RT protected from light. Nuclear staining was performed with 300 nM DAPI in PBS-T for 10 min at RT. Finally, the cells were washed 2x with PBS-T (10 min per wash) followed by 3x washes with DPBS. The coverslips were mounted on microscope slides using Fluoromount-G (Thermo-Fisher, 00-4958-02) and imaged using EVOS M5000 fluorescence microscope with the 10x or 20x objective.

### Karyotyping

8 ul of Karyomax Colcemid solution 10 ug/mL in PBS (Gibco, 15212-012) was added to two 35 mm dishes, with cells in logarithmic growth phase at about 70% confluence and incubated at +37°C and 5% CO_2_ for 4 hours. Cells were then dissociated with 600 µl Accutase to get a single-cell suspension and transferred to a centrifuge tube with 1 mL E8 medium and centrifuged at 214 g for 5 minutes. The supernatant was discarded, and cells resuspended in the small volume left by tapping. 3 mL of hypotonic KCl solution was added while gently vortexing and incubated at 37°C for 10 minutes. 3 mL of 3:1 methanol:acetic acid was added drop by drop to cells while gently vortexing, then centrifuged at 214 g for 10 minutes. Supernatant was discarded and cells resuspended in small volume left by tapping, and the process of fixative addition and resuspension repeated twice more. Cells were stored in 1 mL fixative at +4°C before G-banding service was performed by Ambar Labs, Barcelona.

### Overexpression plasmids

To investigate the functional consequences of the *INTS6* protein-truncating variant, wild-type and mutant *INTS6* alleles were overexpressed in HEK293T cells to assess protein truncation. Total RNA was extracted from a patient-derived iPSC line 1494 and reverse-transcribed using the ProtoScript cDNA kit (New England Biolabs, NEB). Full-length INTS6 cDNA was amplified using Q5 High-Fidelity DNA Polymerase (NEB) with gene-specific primers (forward: 5’-GTCCCCGGCCAGCACTATG-3’; reverse: 5’-ACACGCAAGAGTCAATGACAAGC-3’), and restriction sites for XhoI or NheI (for pCMV or pAAV vectors, respectively) and EcoRI were added to the forward and reverse primers. PCR products were cloned into the pCMV-MYC expression vector (Promega) linearized with EcoRI and XhoI (NEB) and transformed into DH5α E. coli. Plasmids were extracted from 5 mL LB cultures using the NucleoSpin Plasmid Mini kit (Macherey-Nagel, 740588.50), and Sanger sequencing was used to select two wild type and two mutant clones. Plasmid DNA (2 µg) was transfected into HEK293T cells (6 × 10⁵ cells/well in 6-well plates) using 6 µL FuGene 6 (Promega) in 600 µL of DMEM High Glucose (EuroClone) supplemented with 10% fetal bovine serum (Gibco), 1% L-glutamine (BioWhittaker), and 1% penicillin/streptomycin (BioWhittaker). An empty vector served as a negative control. Cells were incubated at +37°C with 5% CO₂, and the samples were collected that protein was extracted 36 hours post-transfection.

### Protein extraction

Cell pellets of 1×10^6^ cells were lysed with 100 µL RIPA-buffer (Cell Signaling, 9806) supplemented with 10 µL/ml of Halt™ protease inhibitor (Thermo Fisher Scientific, 10137963). Cell lysates were incubated on ice for 30 minutes, centrifuged at 12000 rcf for 10 minutes at +4°C, and supernatant collected. Protein concentration was quantified with the Pierce™ BCA Protein Assay Kit (Thermo Scientific, 23225) prior to Western blotting.

### Western blotting of INTS6 and RNAPII pSer2

5-20 µg of protein sample was added to 1:3 of the total volume 4 x Laemmli loading buffer (BioRad, 1610747) containing 10 % β-mercaptoethanol (BioRad, 1610710). Samples were loaded on 4-20% gradient Mini-PROTEAN-TGX SDS-PAGE gels (BioRad, 456-8033), with Precision Plus Protein™ Standard (BioRad, #1610374) as the size ladder. SDS-PAGE were transferred onto a 0.45 µM nitrocellulose membrane (BioRad, 1620234) by semi-dry blotting using the Trans-Blot Turbo transfer system (BioRad). The membranes were blocked in 5 % milk (Valio) in TBS with Tween 20 (0.1 %), or in 5% BSA/TBS-T. The primary antibody for INTS6 was anti-DICE1 antibody mouse (Santa-Cruz, sc-376524) and was diluted in 1% milk/TBS-T. Primary antibodies diluted in 1% BSA/TBS-T were Phospho-Rpb1 CTD (Ser2) (E1Z3G) Rabbit mAb (Cell Signaling Technology, #13499) and Rpb1 NTD (D8L4Y) Rabbit mAb (Cell Signaling Technology, #14958). Loading controls were anti-β-Actin antibody, mouse monoclonal (Sigma-Aldrich, A1978-100UL), or anti-β III-tubulin antibody (rabbit) (Sigma, T2200-200UL). Secondary antibodies were IRDye® 800CW Goat anti-Mouse IgG Secondary Antibody (LI-COR, 926-32210) and IRDye® 680RD Goat anti-Rabbit IgG Secondary Antibody (LI-COR, 926-68071). Images were detected with Odyssey Clx Imaging System v. 6.0.0.28, and quantification of band intensity relative to housekeeping protein was done with ImageStudio software v. 6.1.0.79 (LI-COR). Blots were performed on three independent biological replicates for case and control iPSCs and at least three technical replicates per line. Western blot measurements from repeated gels were treated as repeated technical measurements of the same biological samples. Group differences were analysed using nested ANOVA and pairwise comparisons tested using Tukey-adjusted post-hoc tests for case-control comparisons, and using a linear mixed-effects model with genotype as a fixed effect and gel as a random effect, with pairwise comparisons tested using estimated marginal means and Tukey-adjusted post-hoc tests, for comparisons involving CRISPR correction, het KI, and hom KI (KO) lines. All analyses and visualizations were conducted in RStudio (v. 2025.9.0.387) and ggplot2 v.4.0.2^69^.

### Bulk RNA-sequencing

Total RNA from case and control cell pellets at three timepoints of neuron differentiation (6×10^5^ – 1×10^6^ cells/pellet) were extracted using the NucleoSpin RNA kit (Macherey-Nagel, 740955.250) with rDNAse treatment according to the manufacturer’s instructions.

Transcriptomic library preparation with Illumina Stranded Total RNA Prep with Ribo-Zero Plus, MiSeq Nano v2 2×150bp for sample quality control, and sequencing with NovaSeq S2 200 c, 2×100bp with an average of 73.2 million sequence reads/sample for cases and controls and 239 million reads/sample for CRISPR-engineered samples was provided by the Institute for Molecular Medicine Finland FIMM Genomics unit.

### CRISPR-Cas9 genome editing

To engineer specific genetic modifications in the *INTS6* gene, we targeted the chr13:51378376:G:A variant to knock-in into control iPSC line HEL252.2 and to correct it in case iPSC line HEL246.2. Two independent single guide RNAs (sgRNAs) were designed for both the knock-in and isogenic corrected edits based on GRCh38 using the web tool Benchling (https://benchling.com/crispr). The sgRNAs were designed to be 20 nt, with Cas9 protospacer adjacent motif (PAM) NGG. gRNAs were picked based on predicted on-target and off-target scores. The specificity and efficiency of the chosen sgRNAs was tested *in silico* with CRISPOR version 5.0.1 (http://crispor.tefor.net/), with cutting frequency determination (CFD) scores >60 considered sufficient for genome editing. 101 nt long homology directed repair (HDR) templates for each gRNA were designed using Benchling (gRNAs and HDR templates described in **Table S21**). The sgRNAs and ssDNA HDR templates were supplied by Integrated DNA Technologies (IDT). For each edit, the corresponding sgRNA, ssDNA repair template and HiFi Cas9 were electroporated into iPSC using the Lonza Nucleofector in two replicates per edit. sgRNA activity was measured at the bulk level using T7E1 assay, after which clones were created by low-density seeding and manual colony picking and divided into duplicate 96-well plates, one for genotyping and one for freezing and downstream applications. Genotyping of clonal cell lines was done using MiSeq 150bp paired-end sequencing at the Institute for Molecular Medicine Finland FIMM Genomics unit. The clonal cell lines with desired edits based on MiSeq(N=2 each for KI, KO, and isogenic correction) were expanded and re-genotyped using Oxford Nanopore sequencing to ensure clonality, followed by Sanger sequencing. The clones with the confirmed edits were screened for off-target editing in three top-off targets identified with CRISPOR^70^, and characterised with immunocytochemistry for markers of the undifferentiated state and karyotyped as previously described. CRISPR lines are described in **Table S15.**

Expanded methodology and QC analyses for the CRISPR editing and clone screening are described in a methods manuscript (Forsen *et al*, in preparation) from the Kilpinen lab.

### Sanger sequencing for on- and off-target edits in CRISPR-edited clones

Genomic DNA was extracted from iPSC lines according to manufacturer’s instructions using the NucleoSpin Tissue kit, Macherey Nagel, 740952.50. A 10 ul PCR reaction per clone was set up using 500 bp primers (Sigma) around the *INTS6* chr13:51378376:G:A variant site in CRISPR-edited clones and their respective parental iPSC lines. The same PCR reactions were done for the CRISPR clones and parental lines for the three top off-targets identified using CRISPOR. PCR products were cleaned using ExoSAP-IT PCR Product Cleanup Reagent (Thermo Fisher Scientific, 78201) following the manufacturer’s instructions. For each sequencing reaction, 5 μl of cleaned PCR product at a concentration of 5 ng/μl was mixed with 5 μl of the forward sequencing primer (5 pmol/μl) prepared in nuclease-free water. Premixed samples (final volume ≥10 μl) were transferred into Eurofins Mix2Seq tubes, sealed with the provided caps and submitted for sequencing at Eurofins Genomics. Electropherograms were inspected manually, and sequences aligned to an Hg38 reference sequence using SnapGene v 8.2.1 and NCBI-BLAST. Off-target editing effects were identified by comparing the region spanning ±30 bp around the predicted Cas9 cut site. Primers listed in **TableS22**. On-target *INTS6* electropherograms are in **Fig. S7A,** off-target electropherograms in **Fig. S7B**, and off-target genotyping results described in **Table S23.**

### Northern blotting

For Northern blotting, Trizol-extracted total RNA (1 µg) or IP RNA was run on a 7% urea-polyacrylamide gel, transferred onto Zeta-Probe (Bio-Rad) or Hybond N+ (Cytiva) nylon membrane using an Owl VEP-2 blotter and crosslinked using UV light. Membranes were probed with [γ-^32^P]-ATP-labeled DNA or LNA/DNA oligonucleotides listed in **TableS19**. Hybridization was carried out overnight at 42°C (DNA) or 45°C (LNA/DNA) in hybridization buffer (6xSSC, 25 mM Na_2_HPO_4_/NaH_2_PO_4_ (pH 7.4), 0.5% SDS, 5× Denhardt’s solution, 150 μg/ml yeast RNA (Roche), additionally containing 50% formamide for DNA/LNA probes. Membranes were washed at room temperature with 2x SSC, 0.1% SDS and 0.5x SSC, 0.1% SDS, followed by additional washes at room temperature and 60°C with 0.1x SSC, 0.1% SDS for LNA/DNA probes, 15 min for each wash. Blots were exposed on imaging plates which were scanned using the Sapphire FL Biomolecular Imager.

### Transcriptomic analyses

Sequencing data were processed using the ePouta computing environment for sensitive data provided by the Finnish IT Center for Science Ltd. (CSC) and University of Helsinki. Sequencing quality control was done using FastQC^71^. Sequences were aligned against mixed human and rat reference genomes GRCh38.109 and mRatBN7.2.109 using STAR v.2.7.10a^72^ The mouse *Ngn2* gene was included in the reference to estimate expression of the transgene. Sequence counts were summarized from exons to genes with Subread featureCounts (options -p -t exon -g gene_id -s 2)^73^ keeping uniquely mapped reads. Sequencing alignment summary statistics described in **Table S2** for case and control iPSC- and iPSC-derived NPCs and neurons, **and Table S15** for CRISPR-Cas9 edited iPSCs).

Differential gene expression analysis was performed with limma v. 3.60.1^72^. Raw count data was normalized with the trimmed mean of M-values (TMM), followed by voom transformation prior to linear modelling. The main sources of variation and data dimensionality were assessed using the R packages variancePartition^74^ and PCAtools^75^. Based on the variancePartition analysis, sample_timepoint, sex, *Ngn2* transgene expression and the percentage of astrocytes in the culture as cofactors in the linear model **(Fig. S3A-C).**

Technical replicates were modelled as a random effect using the duplicateCorrelation function in the limma package. Gene Set Enrichment Analysis (GSEA) was performed using clusterProfiler v.4.16.0^76^. For visualisation, top-ranked GO Biological Process terms were selected independently for each timepoint (D0, D4, D28) by ranking gseGO() results by adjusted p-value (Benjamini-Hochberg) and retaining the top 10 per timepoint. Redundant terms were then collapsed using semantic similarity (Wang’s method, implemented in GOSemSim v.2.34.0 ^77^; similarity cutoff = 0.7), retaining the most significant representative term from each similarity cluster. The term with the highest significance was selected as the representative term for each cluster to facilitate biological interpretation.

To estimate the level of expression of the mutant allele in cases, we analysed the sequence reads with ASEReadCounter implemented in GATK v4.5.0.0^78^. Prior variant calling alignment files were marked for duplicate reads, spliced reads were split, and mismatching overhangs were hard clipped with markDuplicates and SplitNCigarReads tools. ASEReadCounter was employed to call biallelic SNPs requiring coverage of at least 10 bases of reference or alternative base with minimum quality of 20 in Phred scale.

To quantify differential intron retention between cases and controls, we analysed the RNA-seq data with IRFInder^79^ v2.0-beta, which calculates changes in the percentage of spliced-in (i.e. retained) introns (ΔPSI) and its significance. We used default options except defining wl=4 (i.e. discarding LowCover, LowSplicing, MinorIsoform and NonUniformIntronCover) (Results in **Table S13** and **Table S17**).

To analyse differential splicing between cases and controls we used LeafCutter^80^ Command “Regtools junctions extract” was run with default options “-a 8 -m 50 -M 500000”. When clustering introns, we required a minimum of 50 split reads supporting each cluster in D0 iPSCs and D4 NPCs, and 20 split reads for D28 neurons, given less sequence coverage was available due to co-culturing with astrocytes.

Statistical analyses were done using base R v.4.5.1. Line art was drawn using R ggplot2 v.4.0.2^69^, and Inkscape v.1.4.2.

### Bulk deconvolution

We deconvoluted bulk RNA-seq profiles from 27 samples collected at D28 of Ngn2 differentiation to estimate individual cell type proportions using CIBERSORTx^81^.

CIBERSORTx input consisted of the raw count matrix of our bulk RNA-seq dataset (28,339 genes x 27 samples) and a published single-cell RNA-seq reference^43^ (18,428 genes x 28,734 cells) matched by differentiation protocol and differentiation day. To provide updated and fine-grained cell type identities for deconvolution, we reannotated the single-cell reference using literature-derived markers across five cell type hierarchies with Snapseed^82^. Snapseed is part of the Human Neural Organoid Cell Atlas Toolbox. For this, gene identifiers from the raw counts matrix were converted from Ensembl IDs to gene symbols, while duplicated or unmapped genes were removed. The single-cell reference dataset was reprocessed using Scanpy^83^ (v1.11.5) in Python (v3.10.). Cells with fewer than 200 detected genes and genes expressed in fewer than 3 cells were filtered out. Raw counts were normalized to 1 million counts per cell and log transformed. Highly variables genes were identified and data was then scaled to zero mean and unit variance with clipping of extreme values (max=10). Principal component analysis (PCA) was performed using the “arpack” implementation and batch effects associated with sample identity were corrected using Harmony v1^84^, followed by UMAP visualization. Cells were clustered using the Leiden algorithm (resolution=1.1). The dataset was then reannotated using the ‘annotate_hierarchy’ function in Snapseed, assigning cell identities to the following cell type hierarchies with corresponding cell numbers (N):

- Excitatory neurons (*non-telencephalic/diencephalic*, N=3,281; *non-telencephalic/diencephalic & hypothalamic,*N=2,535; *non- telencephalic/mesencephalic*, N=4,361; *non-telencephalic/rhombencephalic/cerebellar*, N=1,201; *non-telencephalic/rhombencephalic/pons*, N=8,608)
- Peripheral nervous system (PNS) neurons, N=7,673 Neural progenitors (*diencephalic*, N=11; *outer radial glia*, N=349; *rhombencephalic/cerebellar*, N=482)
- Pericytes (N=233).

After formatting both inputs, we ran CIBERSORTx via Apptainer, using a container image converted from Docker. Access was authenticated using a requested personal access token. Deconvolution was performed in fraction mode, enabling batch correction and estimating relative cell type fractions (standard mode).

We then performed differential abundance to test whether the ID condition was impacting cell type proportion using a non-parametric Wilcoxon rank-sum test (Mann-Whitney U test). For downstream analysis and visualization, fine-grained cell type annotations were collapsed into four major categories (excitatory neurons, PNS neurons, neural progenitors and pericytes). Due to their low abundance, pericytes were excluded from subsequent analysis.

## Acknowledgements

The authors would like to warmly thank the study participants of NFID for their willingness to participate in the study. The authors acknowledge funding from Jane and Aatos Erkko Foundation (O.P.), Päivikki and Sakari Sohlberg Foundation (O.P.), Jenny and Antti Wihuri Foundation (O.P.), Instrumentarium Research Foundation (O.P.), Jalmari and Rauha Ahokas Foundation (O.P.), Research Council of Finland (#338835 to H.K.), Sigrid Jusélius Foundation (H.K., M.J.F., P.P.), Helsinki Institute of Life Science, University of Helsinki (H.K.). O.P. and H.K. were additionally supported by Understanding the Human Brain (UHBRAIN) PROFI6 funding from the Research Council of Finland to the University of Helsinki. The reprogramming of iPSC lines was provided by Harvard Stem Cell Institute, Cambridge, MA, USA, and Biomedicum Stem Cell Center, supported by HiLIFE and the Faculty of Medicine, University of Helsinki, and Biocenter Finland. RNA-sequencing was performed at the Institute for Molecular Medicine Finland FIMM Genomics unit supported by HiLIFE and Biocenter Finland. The authors wish to acknowledge CSC – IT Center for Science, Finland, for computational resources.

## Author contributions

Conceptualization: O.P.

Methodology: N.J., K.T., O.P., H.K., M.J.F.

Investigation: N.J., K.T., A.J.N., P.L., L.I., N.N.

Formal analysis: N.J., K.T., P.P., V.V., N.A.

Visualization: N.J., K.T., P.P.

Writing – original draft: N.J., O.P., H.K.

Writing – review & editing: All authors Supervision: O.P., H.K.

Project administration: O.P., H.K., E.H.

Funding acquisition: O.P., H.K., M.J.F.

Resources: L.U., M.K., A.P., O.K., E.R., E.H., O.P.

## Data and code availability

The code generated during this study are available at https://github.com/nelliaj/NFID_INTS6_ID_iPSC_modeling.git

The RNA-seq count data will be deposited in the European Genome-Phenome Archive (https://ega-archive.org/).

