## Supplemental_figures for "*INTS6* loss of function disrupts transcriptional regulation in mild intellectual disability"

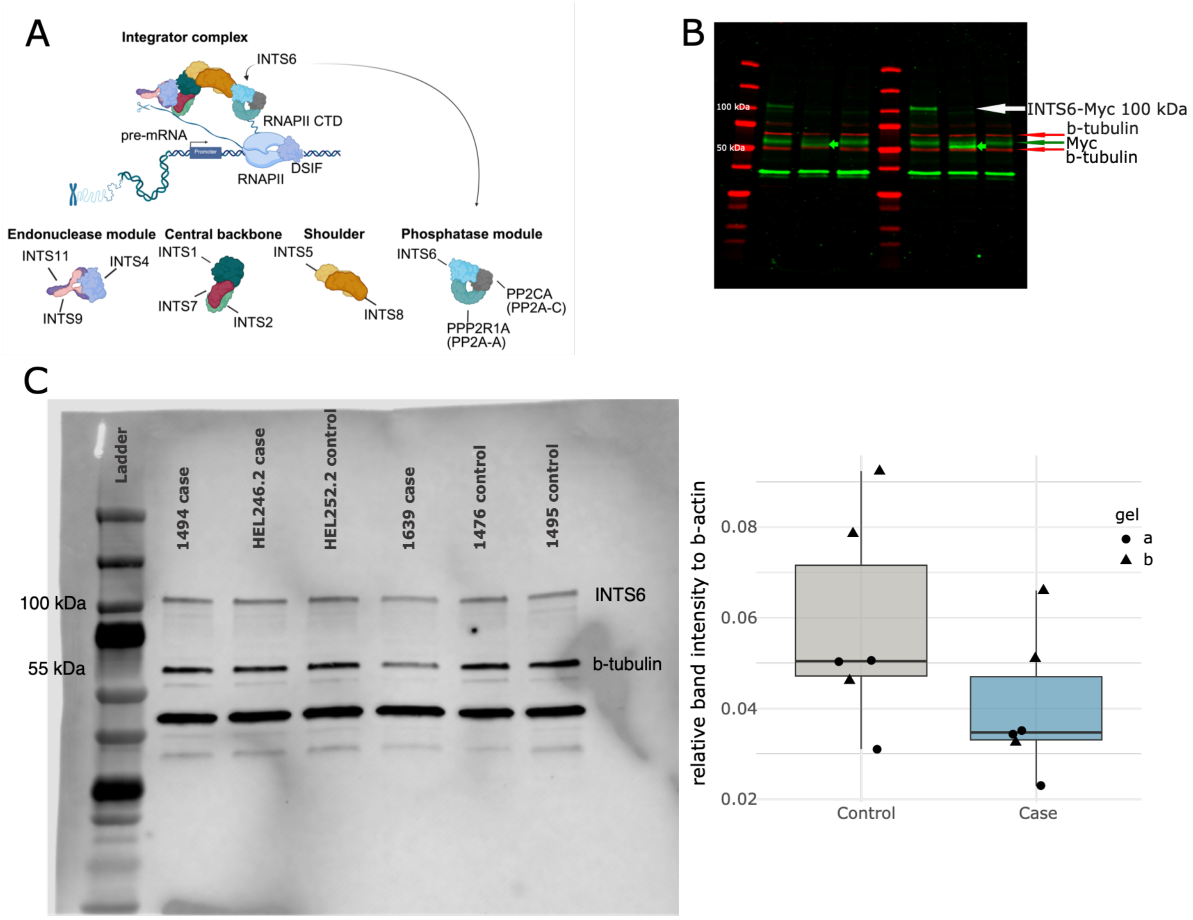


**Figure S1: Characterisation of the protein consequences of *INTS6* c.1465C>T, p.Arg489*. (A)** Schematic of the Integrator complex showing *INTS6* as part of the phosphatase module. Figure adapted from: Welsh, S.A. & Gardini, A. Genomic regulation of transcription and RNA processing by the multitasking Integrator complex. *Nature Reviews* **24**(2023). Panel A created with BioRender. **(B)** Uncropped Western blot showing overexpression of the INTS6-Myc fusion protein containing either the control (WT) or mutant (pR489*) allele in HEK293T cells. Overexpression was done in two technical replicates (r1/r2). Empty plasmid was used as negative control. pR489* samples show a lack of INTS6 protein expression, and the green arrow indicates the 54.6 kDa predicted truncated protein that is expressed in the pR489* samples. The additional band below INTS6-Myc 100 kDa and 54.6 kDa corresponds to endogenous Myc in the cells. **(C)** Uncropped Western blot showing INTS6 protein is not significantly affected by the PTV ($N_{cases}$ = 3, $N_{controls}$ = 3, p = 0.0733, nested ANOVA) (left). Blots were done in three independent biological replicates per group in two technical replicates across two gels. Band intensities normalised to b-tubulin. Quantification boxplots (right) show median + 1.5 IQR, with points representing biological replicates and point shapes represent different gels (technical replicates).

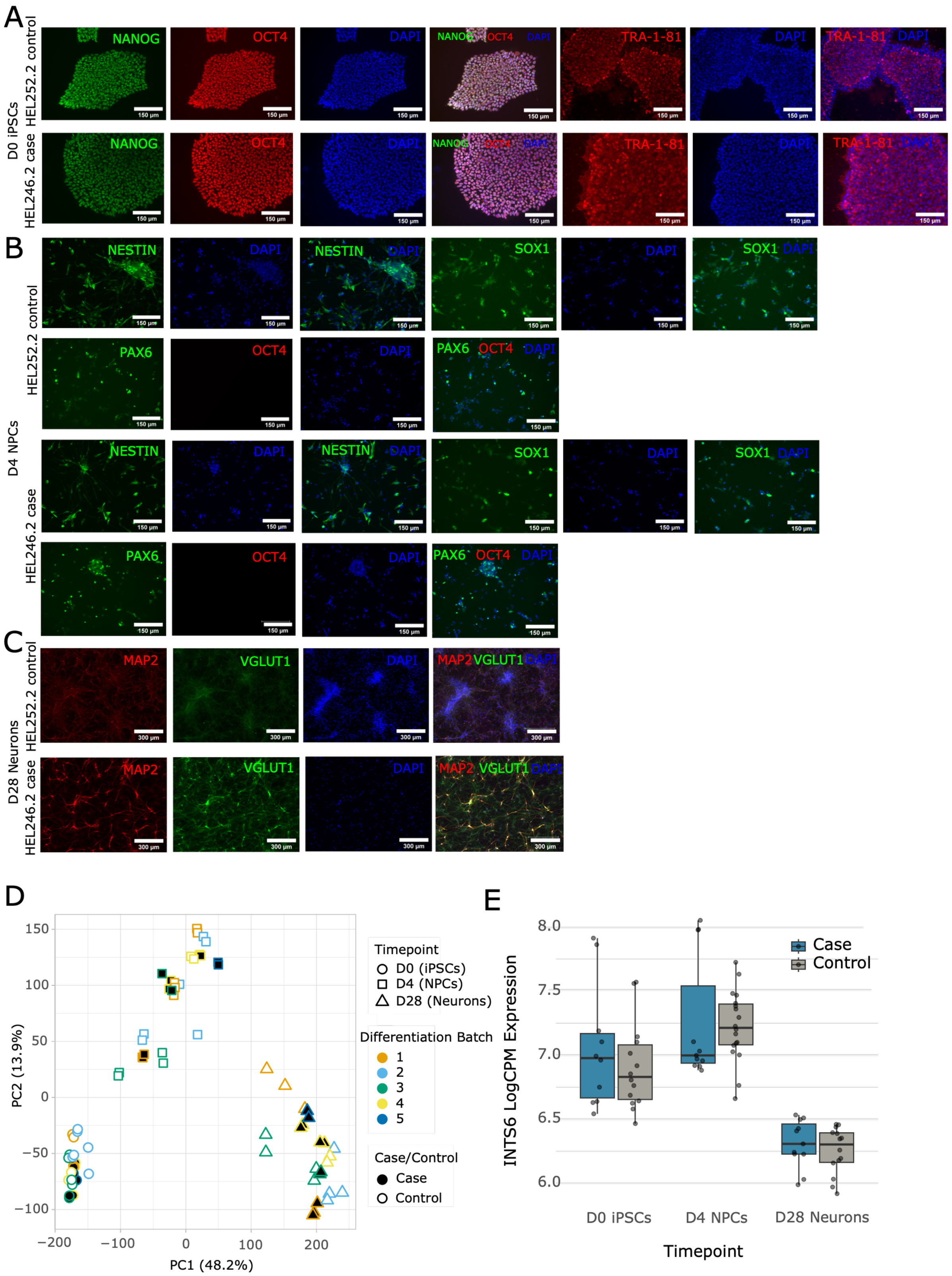


**Figure S2: Characterisation of the iPSCs and iPSC-derived NPCs and neurons from *INTS6* cases and controls. (A)** Immunocytochemistry images (20x) of D0 iPSCs in one representative case and control. **(B)** Immunocytochemistry images (20x) of D4 NPCs in one representative case and control. **(C)** Immunocytochemistry images (10x) of D28 neurons in one representative case and control. **(D)** PCA plot showing PC1 separating differentiation stages, with cases and controls clustering together, and demonstration that differentiation batch (1-5) did not affect differentiation capability. **(E)** *INTS6* mRNA expression was not affected by the *INTS6* LoF variant at any of the differentiation stages (case vs. control, D0 p = 0.283, D4 p = 0.837, D28 p = 0.707, moderated t-test limma-voom).

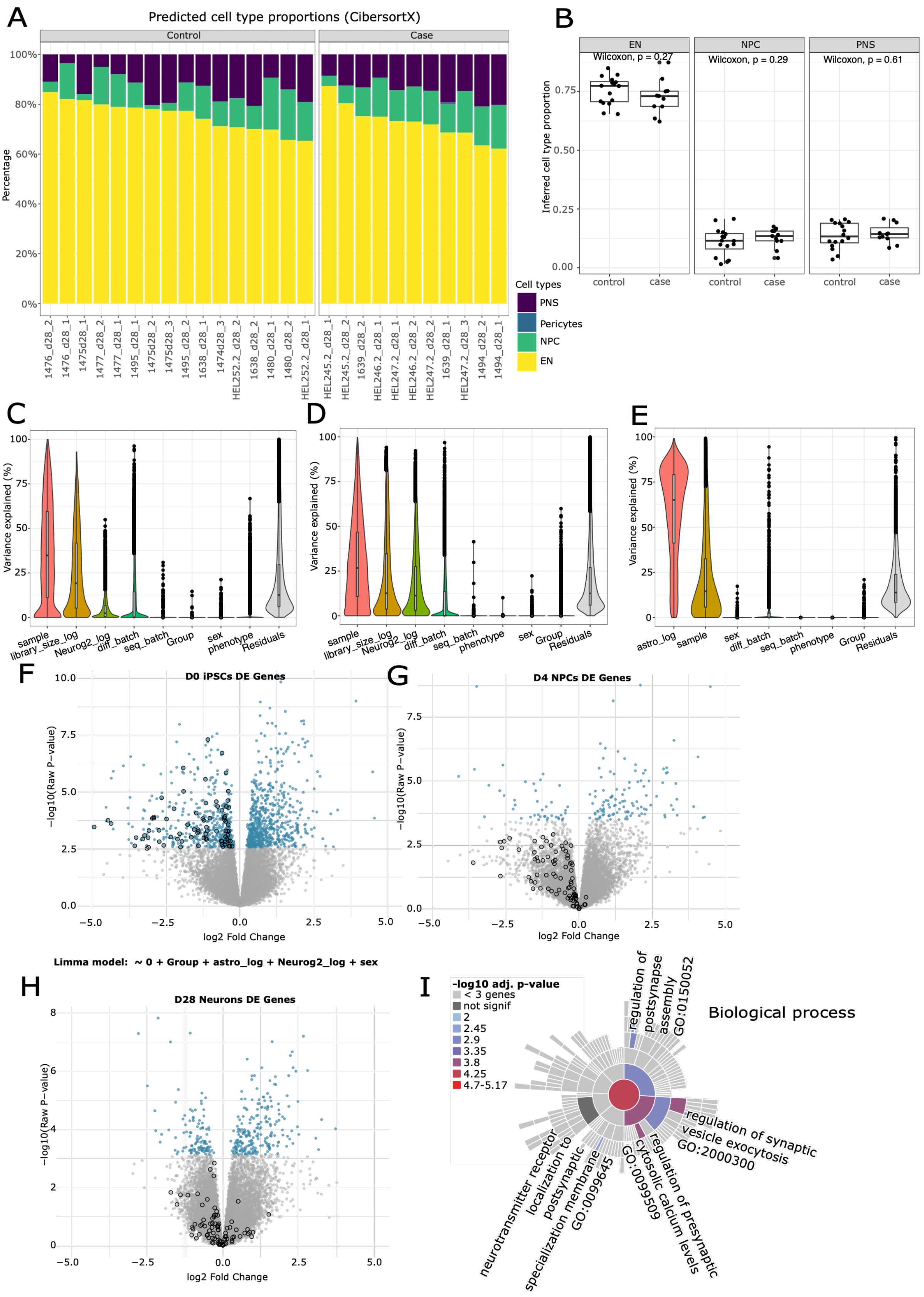
A
Predicted cell type proportions (CibersortX)
Control
Case
Percentage
100%
80%
60%
40%
20%
0%
1476_d28_2
1476_d28_1
1475d28_1
1477_d28_2
1477_d28_1
1495_d28_1
1475d28_2
1475d28_3
1495_d28_2
1638_d28_1
1474d28_3
HEL252.2_d28_2
1638_d28_2
1480_d28_1
1480_d28_2
HEL252.2_d28_1
HEL245.2_d28_1
HEL245.2_d28_2
1639_d28_2
HEL246.2_d28_1
HEL247.2_d28_1
HEL246.2_d28_2
HEL247.2_d28_2
1639_d28_1
HEL247.2_d28_3
1494_d28_2
1494_d28_1
Cell types
PNS
Pericytes
NPC
EN
B
EN
NPC
PNS
Wilcoxon, p = 0.27
Wilcoxon, p = 0.29
Wilcoxon, p = 0.61
Inferred cell type proportion
0.75
0.50
0.25
0.00
control
case
C
D
E
Variance explained (%)
100
75
50
25
0
sample
library_size_log
Neurog2_log
diff_batch
seq_batch
Group
sex
phenotype
Residuals
astro_log
F
D0 iPSCs DE Genes
G
D4 NPCs DE Genes
−log10(Raw P−value)
log2 Fold Change
10.0
7.5
5.0
2.5
0.0
−5.0
−2.5
0.0
2.5
5.0
Limma model: ~ 0 + Group + astro_log + Neurog2_log + sex
H
D28 Neurons DE Genes
8
6
4
2
0
I
-log10 adj. p-value
< 3 genes
not signif
2
2.45
2.9
3.35
3.8
4.25
4.7-5.17
Biological process
regulation of postsynapse assembly GO:0150052
regulation of synaptic vesicle exocytosis GO:2000300
regulation of presynaptic cytosolic calcium levels GO:0099509
neurotransmitter receptor localization to postsynaptic specialization membrane GO:0099645

Fig. S3 contd.

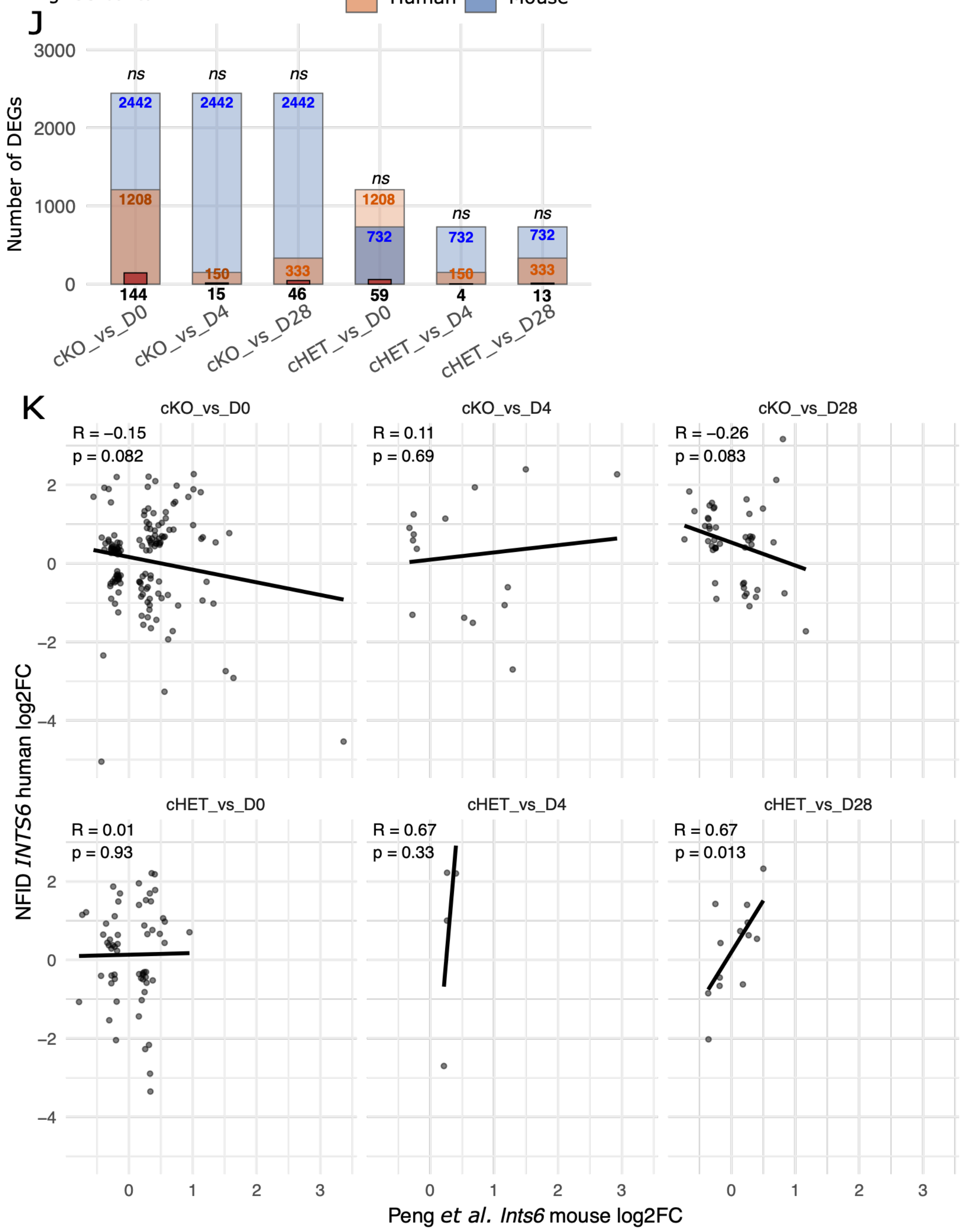


**Figure S3: Exploring the transcriptomic signatures of *INTS6* cases and controls. (A)** Stacked barplot using CiberSortX (v 1.0) deconvolution results showing that percentage of excitatory neurons for all case and control neurons (D28) varied between 62.2-87.4%%. **(B)** Boxplots showing that inferred proportions of excitatory neuron (EN) were not significantly different between cases and controls (p = 0.27, Wilcoxon test), and neither was proportion of NPC (p = 0.29, Wilcoxon test) or PNS (peripheral neurons) (p = 0.61, Wilcoxon test). **(C-E)** Violin plot from variancePartition (v 1.38.1) shows the percentage of variance explained by each covariate for each gene in the RNA-seq data from D0 iPSC (**C**) D4 NPCs (**D**) and D28 neurons (**E**). Most of the RNA-seq variance was explained by sample and (log) library size in D0 iPSC, by (log)library size, and (log)Ngn2 expression in D4 NPC and by (log)astrocyte percentage and sample in D28 neurons. **(F-H)** The Limma DEG model was chosen based on the variancePartition results: ~ 0 + Group + astro_log + Neurog2_log + sex, where Group is sample_timepoint. Volcano plots showing DEGs (adj.p<0.05 coloured in blue) between cases and controls at D0, D4 and D28. Human TF genes that were significantly downregulated at D0 are circled in black for all timepoints. **(I)** Sunburst plot of biological process terms from significantly downregulated DEGs in neurons. **(J)** Summary of the comparison between Peng *et al.* (2025) mouse E15.5 cKO and 2-month cHET DEGs and NFID *INTS6* D0, D4 and D28 DEGs. Translucent bars show total DEGs (mouse in orange, human in blue), and solid bar shows shared DEGs. P= hypergeometric test, BH−adjusted (ns = not significant). **(K)** Scatterplots comparing the log2 fold changes (log2FC) of differentially expressed genes (DEGs) in human *INTS6* D0 iPSCs, D4 NPCs and D28 neurons against mouse *Ints6* E15.5 cKO and 2-month cHET DEGs from Peng et al. (2025). The plot highlights a significant correlation (Pearson R=0.67, p=0.013) among 13 shared DEGs between the mouse heterozygous knockouts (cHET) and human D28 neurons, but no significant correlations for the other comparisons (p>0.05).

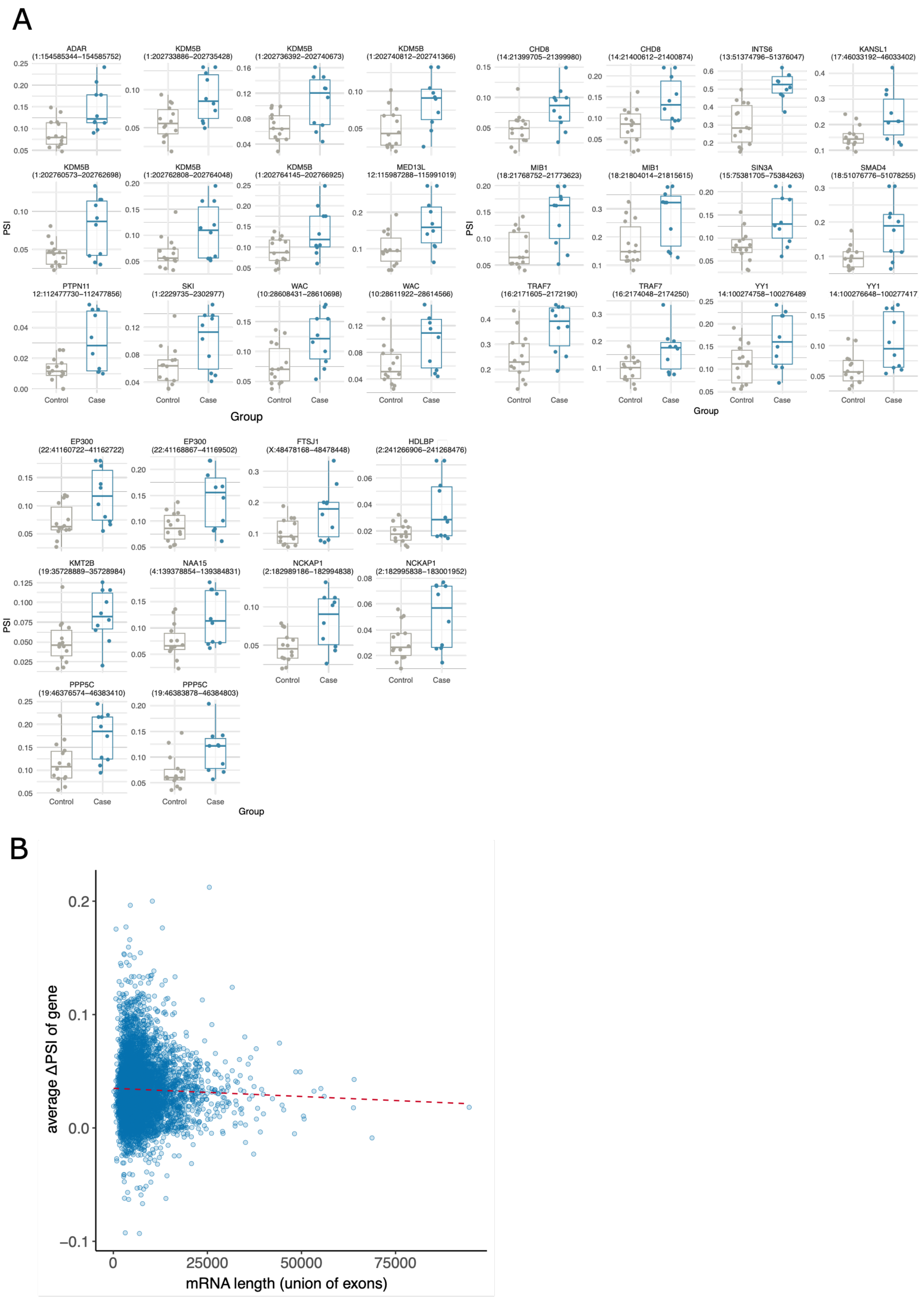


**Figure S4: Intron retention based on IRFinder (A)** Intron retention was significantly higher in *INTS6* cases than controls (ΔPSI> 5%, p.adj <0.05) in multiple introns of 20 known NDD genes: *ADAR, KDM5B, CHD8, INTS6, KANSIL1, MIB1, SIN3A, SMAD4, PTPN11, SKI, WAC, TRAF7, YY1, EP300, FTSJ1, HDLBP, KMT2B, NAA15, NCKAP1*, and *PPP5C*. **(B)**

Correlation between the average ΔPSI of genes with significant IR in cases and their mRNA length defined as the union of its exons. mRNA length was weakly negatively correlated with average ΔPSI of all genes (Pearson R=-0.031, p=0.0253).

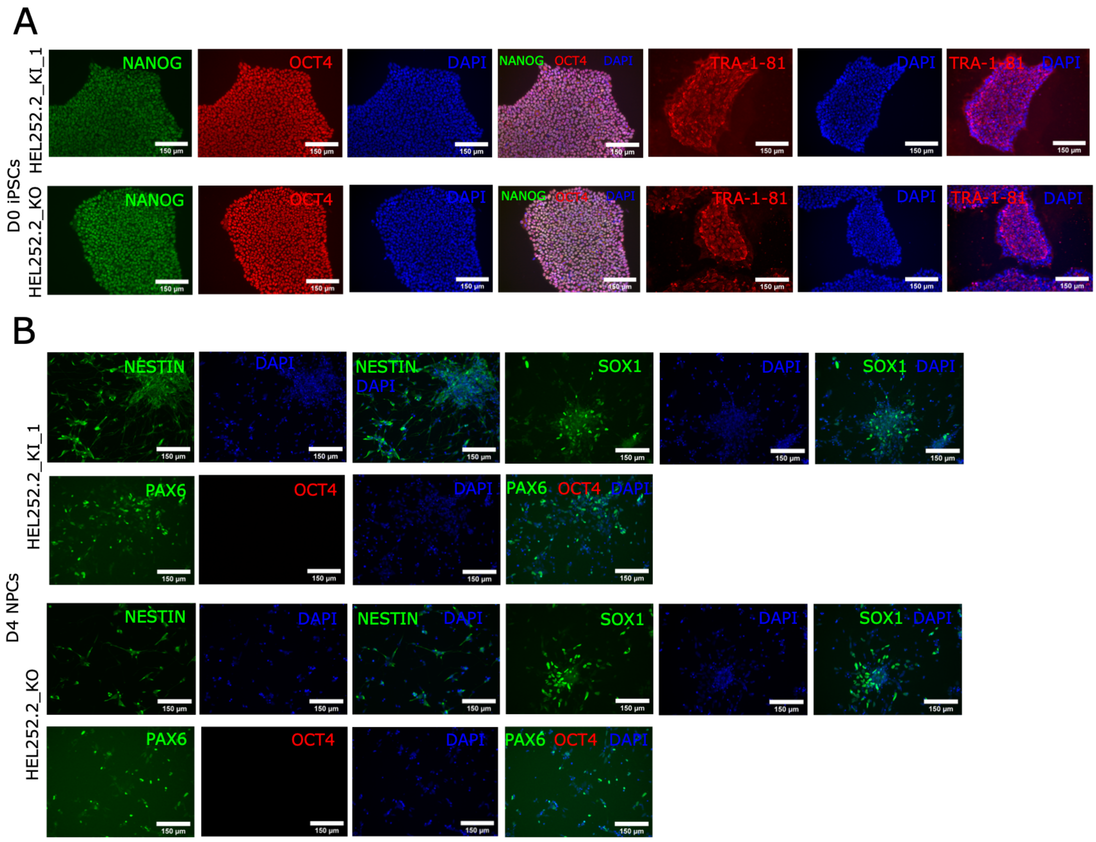

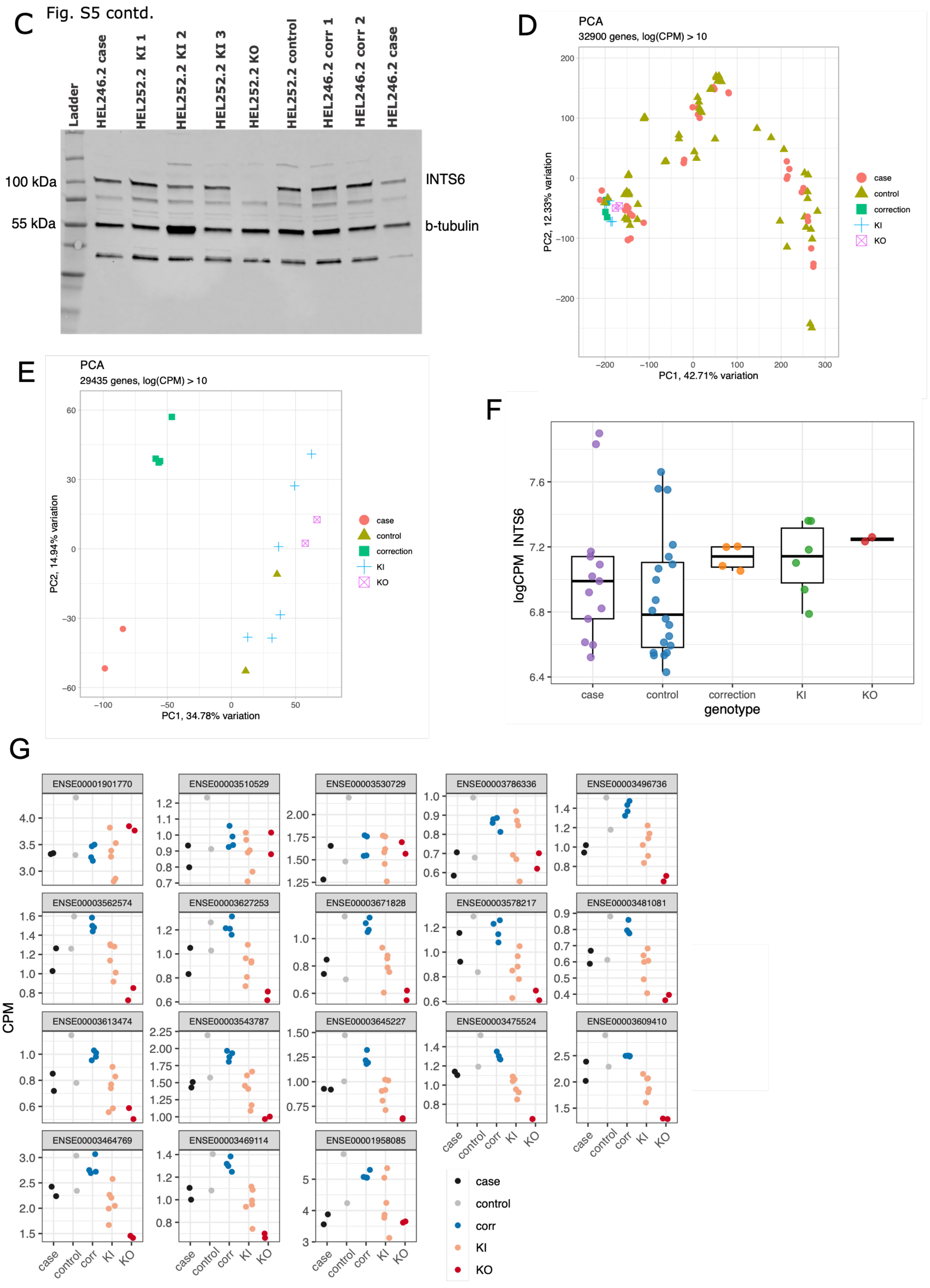


**Figure S5: Studying the transcriptomic and functional effects of the *INTS6* variant using CRISPR-Cas9 edited knock-in and isogenic corrected iPSC lines. (A)** Immunocytochemistry images (20x) of D0 iPSCs in representative CRISPR KI and clones. **(B)** Immunocytochemistry images (20x) of D4 NPCs in representative CRISPR KI and KO clones. **(C)** Uncropped Western blot showing that the KO line produced no detectable *INTS6* protein (100 kDa). **(D)** PCA scatter plot (PC1 vs PC2) showing that, along PC1, CRISPR-Cas9 edited iPSC lines cluster together with the case-control iPSCs. **(E)** PCA scatter plot (PC1 vs PC2) showing that, along PC1, isogenic corrected lines cluster together with their parental case iPSC line and the KI and KO lines cluster together with their parental control iPSC line. **(F)** *INTS6* mRNA expression was unaffected both by the *INTS6* variant of the case, KI and KO lines compared to controls (case-control p = 0.257, KI vs control p = 0.149, KO vs control p = 0.162, all FDR > 0.05 (Benjamini-Hochberg), moderated t-test limma-voom). **(G)** In the CRISPR KO iPSC line, there was a decrease in expression of the exons after the predicted stop-codon, consistent with loss of function.

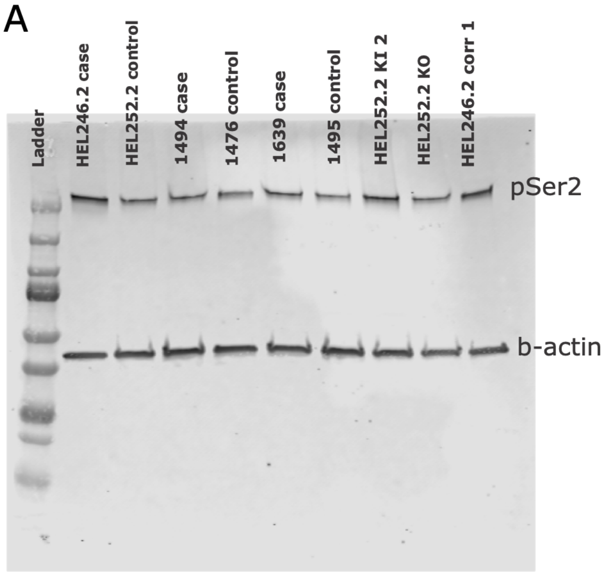


**Figure S6: (A)** Representative uncropped Western blot showing that the *INTS6* LoF variant did not affect RNAPII pSer2 levels.

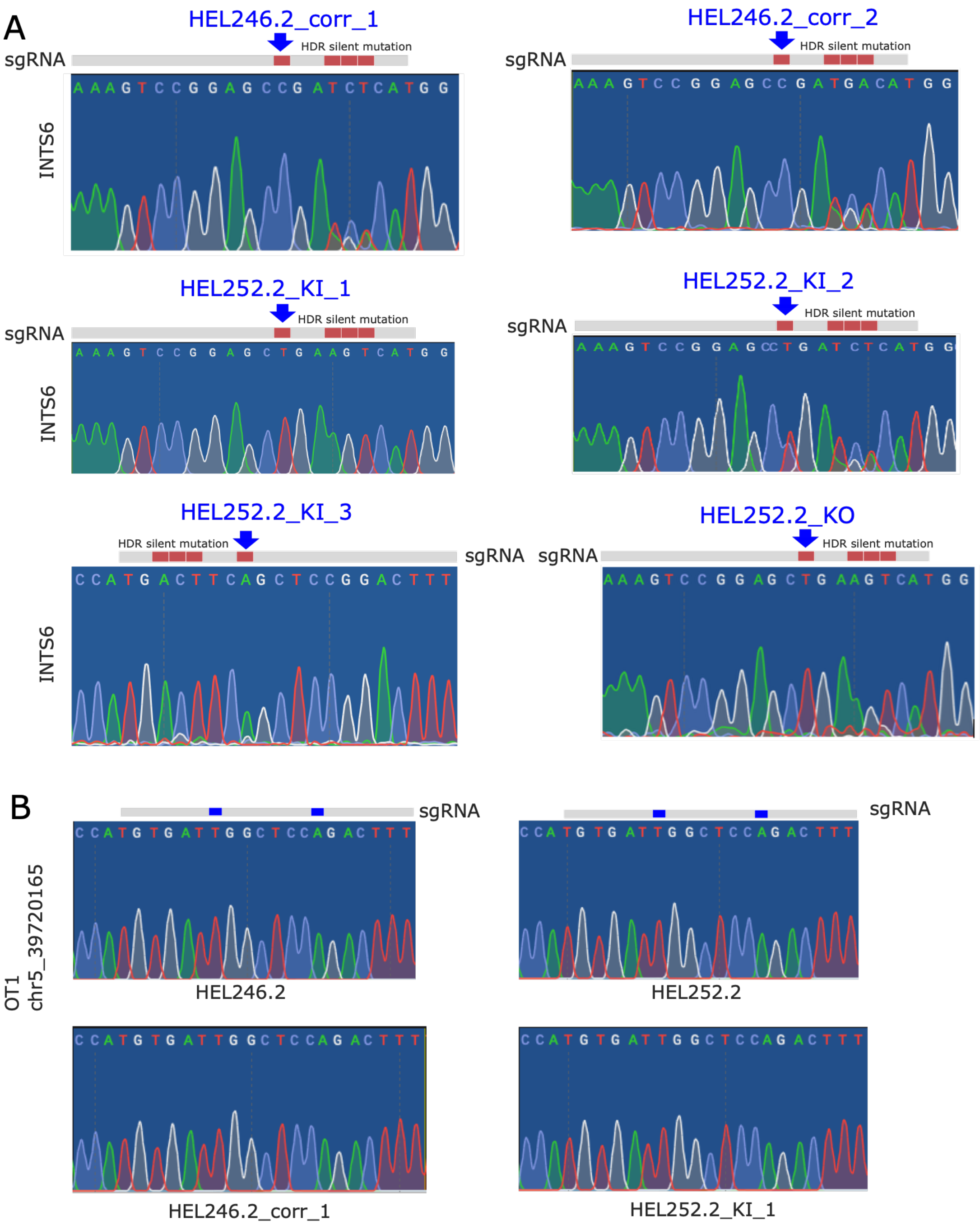


**Figure S7: *INTS6* CRISPR-Cas9 genome editing on-target and off-target genotyping.** **(A)** Sanger sequencing of the region around *INTS6* c.1465C>T showing the CRISPR-Cas9 engineered KI edits made into HEL252.2 and correction edits made into HEL246.2. sgRNA shown in grey and edit site shown by blue arrows. **(B)** Sanger sequencing traces of the top predicted off-target region (OT_1 in chr5_39720165) of the sgRNAs (INTS6_HEL252_RV_sgRNA_1 and INTS6_HEL246_RV_sgRNA_3 in TableS21) as identified by CRISPOR (see Methods). No off-target editing was detected in representative CRISPR correction and KI lines with reference to the parental iPSC lines HEL246.2 and HEL252.2.
